## Supplementary data for "Biallelic and gene-wide genomic substitution for endogenous intron and retroelement mutagenesis in human cells"

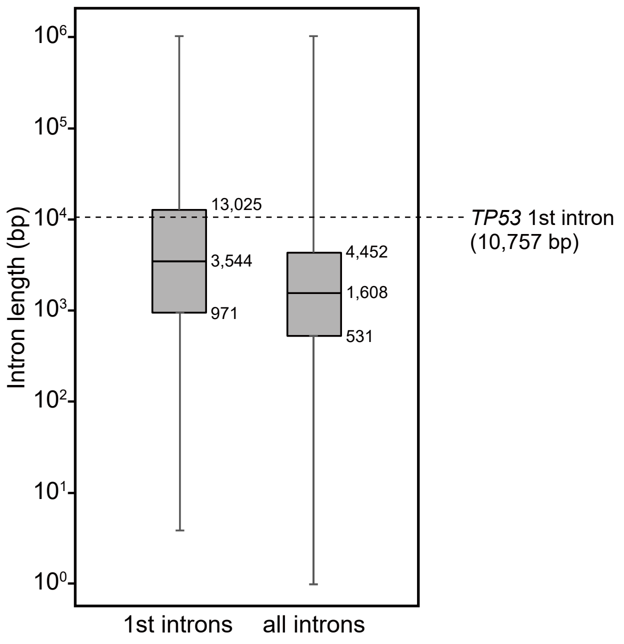


**Supplementary Figure 1 | Box plots represent the distribution of intron length for first introns and all introns within human genes.**

The middle line, the box, and the whiskers indicate the median, the 25th to 75th percentiles, and the interquartile range, respectively. The broken line represents the length of the first intron of *TP53*.

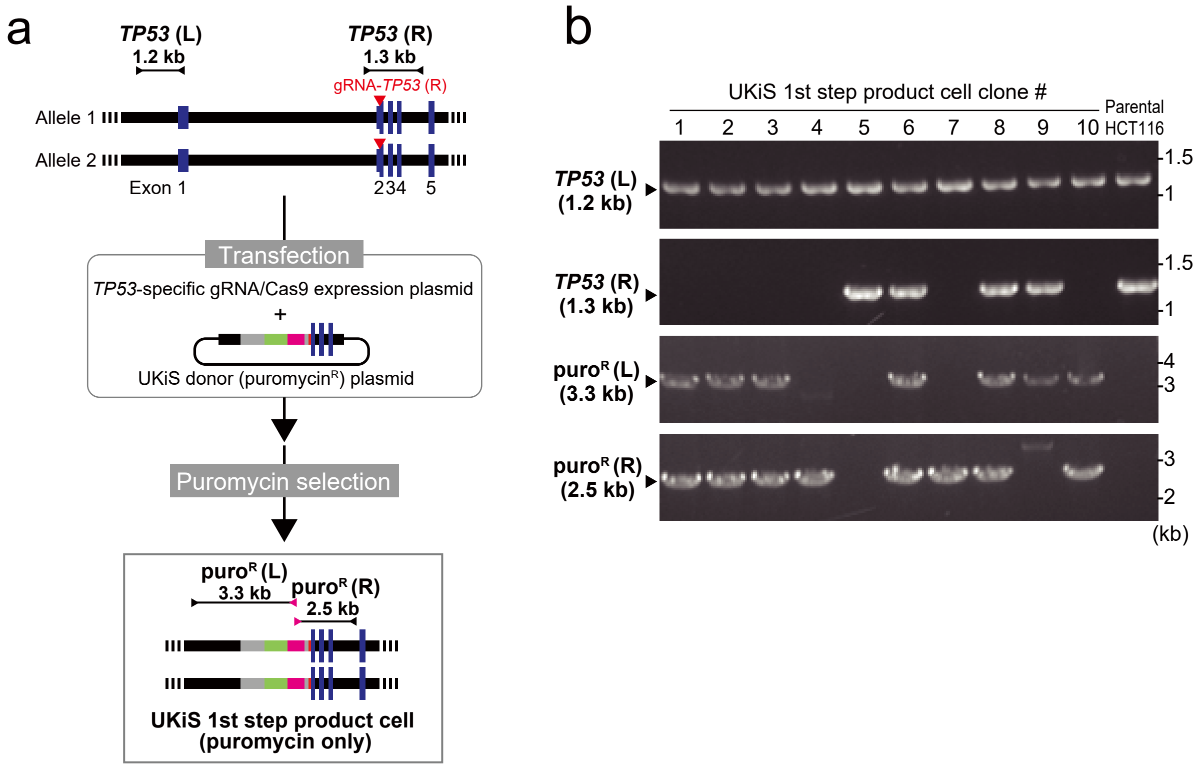


**Supplementary Figure 2 | Replacement of the first intron of *TP53* with only one kind of UKiS donor.**

**a.** Schematic of the UKiS first step for replacement of the *TP53* first intron by using only the puromycin UKiS donor. Horizontal lines flanked by two arrowheads represent the target regions for the junction genotyping PCR.

**b.** Representative gel image of the junction genotyping PCR to confirm deletion of the *TP53* first intron and insertion of the UKiS marker. Of the 10 clones obtained after puromycin selection, none had successful recombination at both alleles.

a

| Off-target candidate | Sequence | Indel | Mismatch | Chr position |
|---|---|---|---|---|
| #1 | CTCAGGGTGGGCTGGACGCTGGG | 0 | 3 | Chr16:89621342-89621364 |
| #2 | CTCAGA-AGGGCTCCACGCTTGG | 1 | 2 | Chr2:191021507-191021528 |

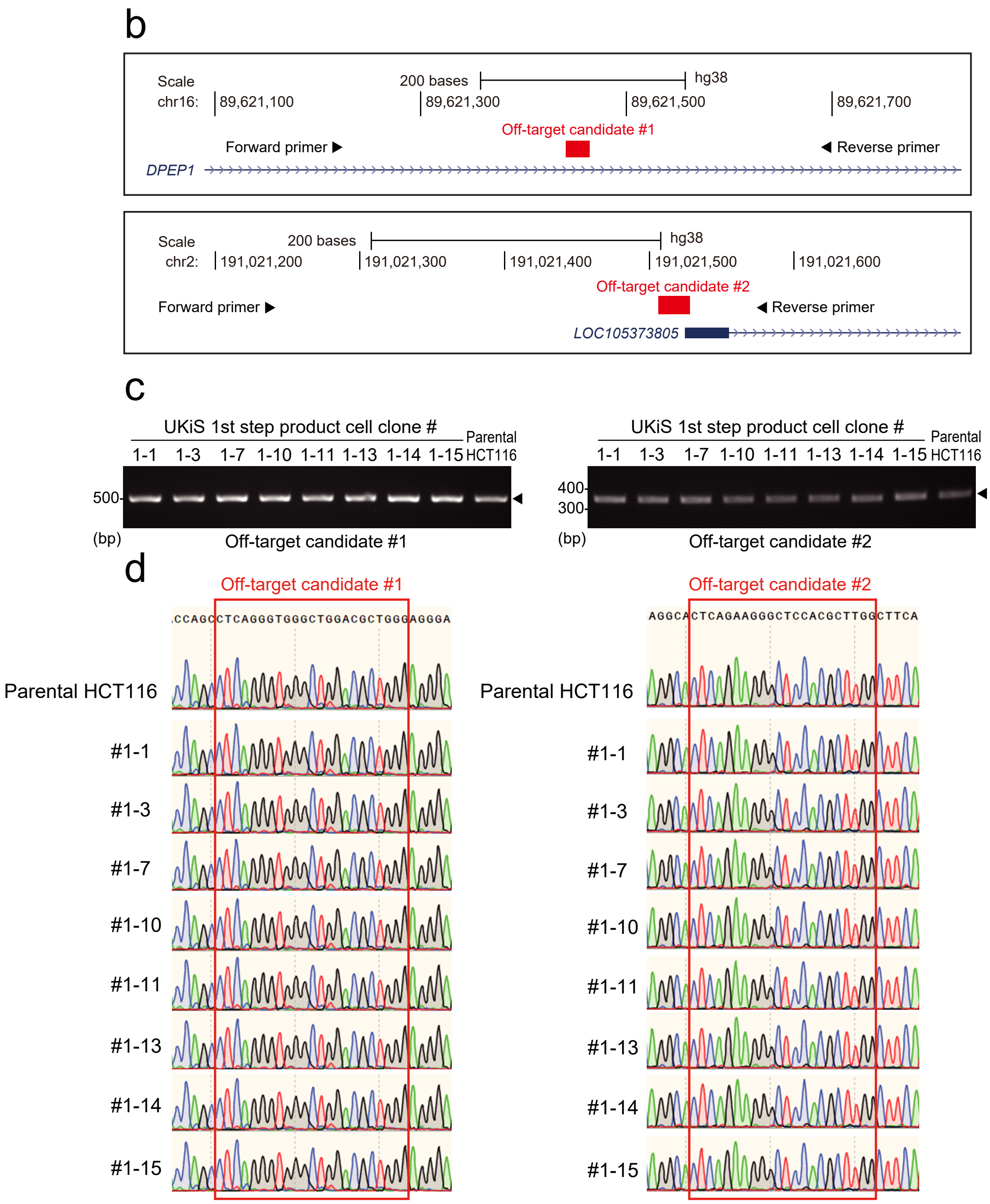


**Supplementary Figure 3 | Assessment of off-target cleavage by the *TP53*-specific gRNA.**

**a.** The two most likely off-target candidate sites predicted by COSMID (https://crispr.bme.gatech.edu/). Mismatched nucleotides and a missing nucleotide relative to the *TP53*-specific gRNA sequence are indicated in red. The PAM sequence is underlined.

**b.** The genomic locations of the off-target candidate sites. The primers used for genomic PCR are denoted by arrowheads. Locations are based on the human genome reference sequence hg38.

**c.** Representative images of genotyping PCR against the off-target sites by using genomic DNA from the indicated cell clones as the template. Genomic DNA was extracted from the positive clones obtained after the first step of UKiS. All clones were confirmed to have undergone successful recombination of the *TP53* intron with the UKiS donors at both alleles (as shown in Fig. 2c).

**d.** Sequencing of PCR amplicons obtained in (c) demonstrated that no indels or any other genomic modifications occurred at either off-target candidate site, indicating that these sites were not mistakenly cleaved by the *TP53*-specific gRNA.

a

| Off-target candidate sequence | Indel | Mismatch | Chr position |
|---|---|---|---|
| GGCGC-AGGCGATCGAGTAAGGG | 1 | 2 | Chr19: 35545383-35545404 |

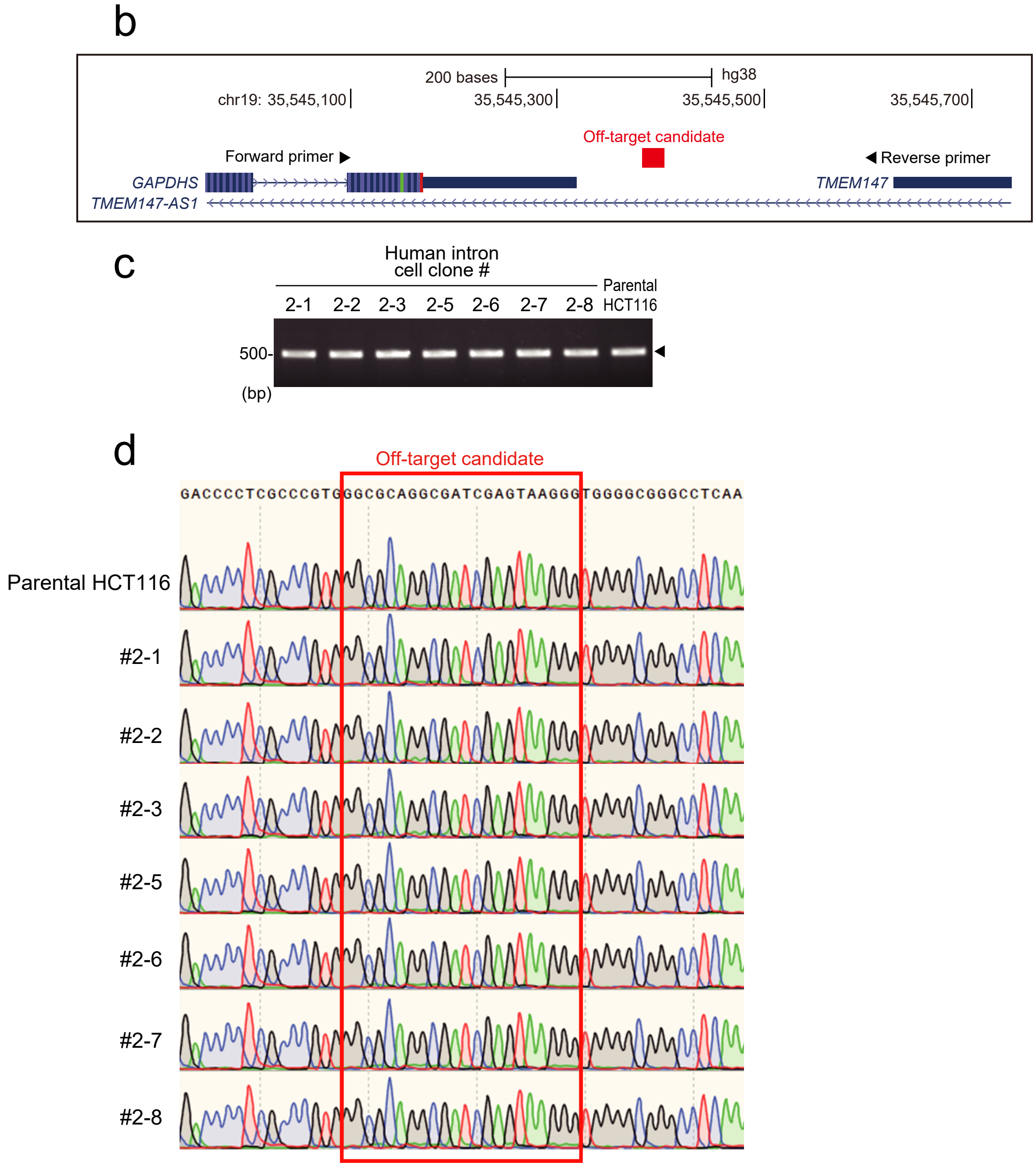


**Supplementary Figure 4 | Assessment of off-target cleavage by TL-gRNA.**

**a.** The most likely off-target candidate site predicted by COSMID (https://crispr.bme.gatech.edu/). Mismatched nucleotides and a missing nucleotide relative to the TL-gRNA sequence are indicated in red. The PAM sequence is underlined.

**b.** The genomic location of the off-target candidate site. The primers used for genomic PCR are denoted by arrowheads.

**c.** Representative image of genotyping PCR against the off-target site by using genomic DNA from the indicated cell clones as the template. Genomic DNA was extracted from the positive clones obtained after the second step of UKiS with the mutating payload containing the wild-type human intron. As shown in Fig. 3b, these clones were all confirmed to have undergone successful recombination of the UKiS donor allele with the human intron.

**d.** Sequencing of PCR amplicons obtained in (c) demonstrated no indels or any other genomic modifications at the off-target candidate site, indicating that this site was not mistakenly cleaved by the TL-gRNA.

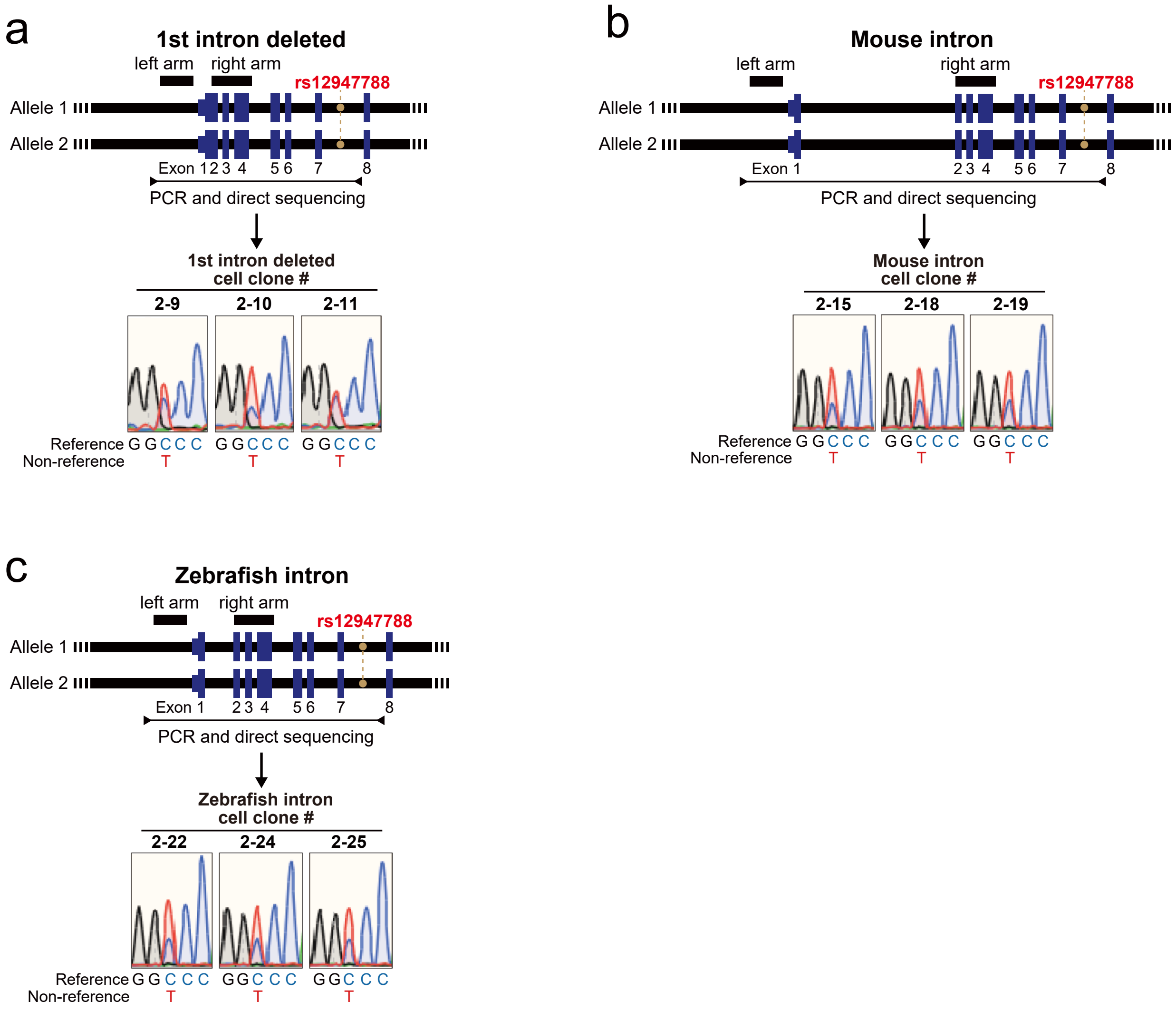


**Supplementary Figure 5 | Validation of biallelic replacement in the second step of UKiS to generate three different kinds of *TP53* first intron mutant clones.**

**a, b, c.** For the (a) intron-deleted clones, (b) mouse intron clones, and (c) zebrafish intron clones, graphical representation of the human *TP53* locus is shown on the top: black boxes represent the positions of the homology arms used in our UKiS mutagenesis to *TP53*, a horizontal line flanked by two arrowheads represents the target region for PCR, and the orange filled circles denote the position of the heterozygous SNP site, rs12947788. Direct sequencing of the PCR genotyping amplicons indicated double peaks only at rs12947788 on the resultant sequencing chromatograms for all three clones for each kind of mutant.

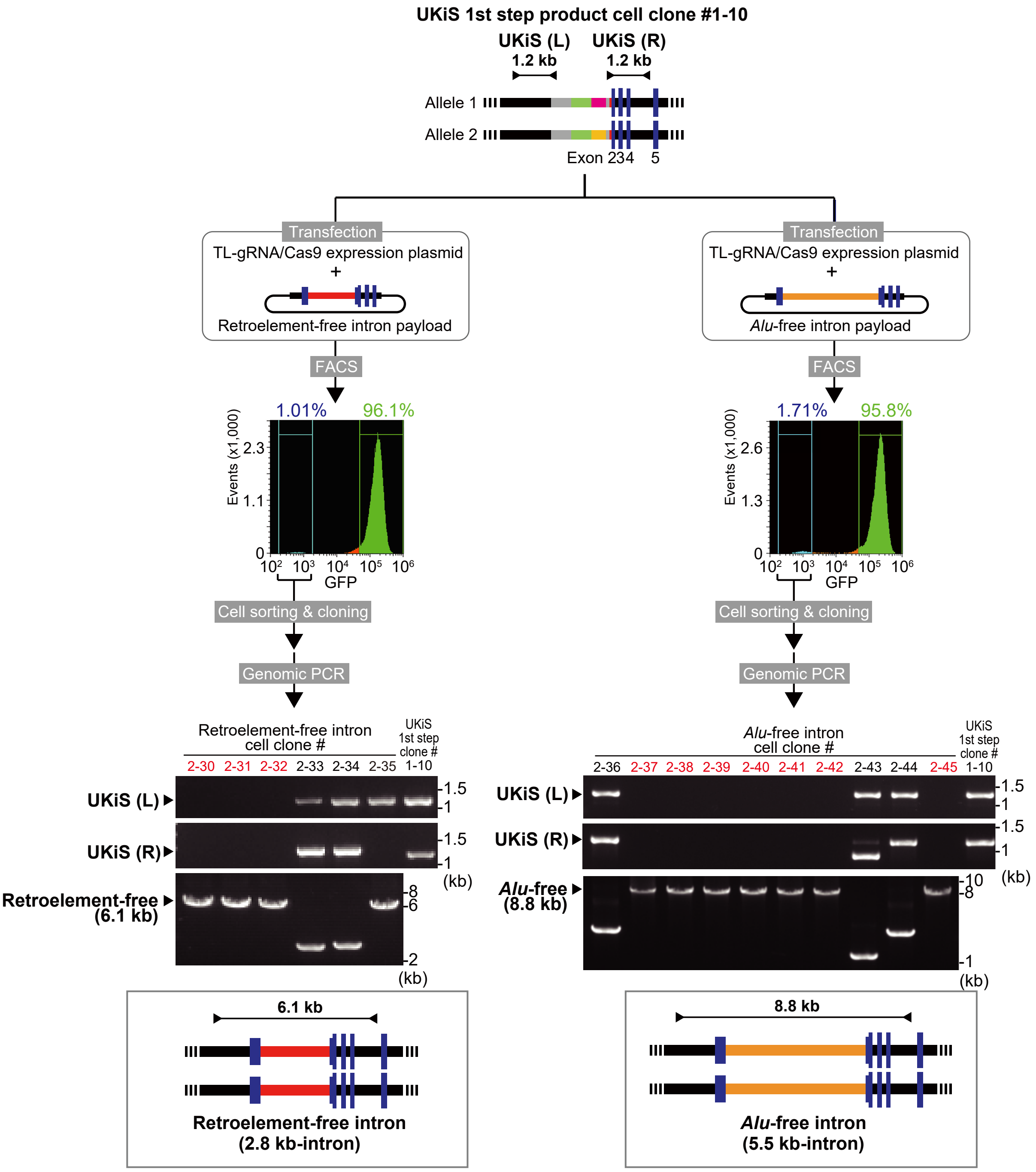


**Supplementary Figure 6 | The second step of UKiS: replacement of both UKiS donor alleles in the first intron of *TP53* with either a retroelement- or *Alu*-free intron.**

Replacement of both UKiS donor alleles in clone #1-10 with one of two synthetic introns from which all retroelement (left) and *Alu*-derived (right) sequences had been removed. First, the mutating payload plasmids containing the retroelement- or *Alu*-free intron and the TL-gRNA/Cas9 expression plasmid were transfected into clone #1-10. Thereafter, GFP-negative cells were collected by FACS and cloned. Biallelic substitution of UKiS markers with the mutating payload plasmid was confirmed by junction genotyping PCR that targeted the regions represented by horizontal lines flanked by arrows in the schematic diagrams of the *TP53* locus after successful replacement, with the expected length of PCR genotyping amplicons indicated. In the agarose gel images, lane numbers of clones that underwent successful recombination in both alleles are in red.

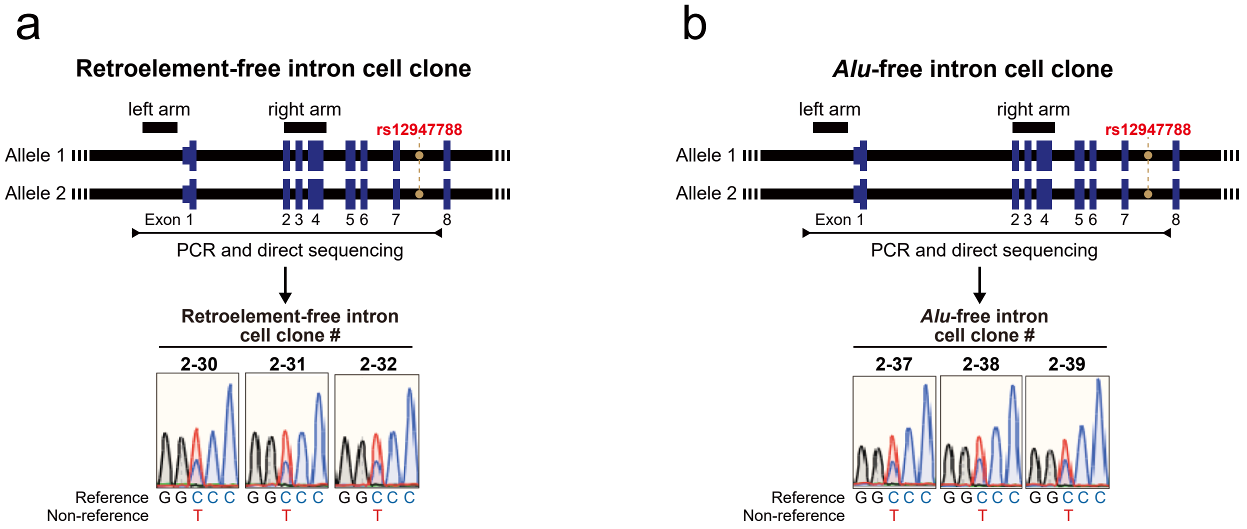


**Supplementary Figure 7 | Validation of biallelic replacement in the second step of UKiS for making the retroelement- and *Alu*-free intron clones.**

**a, b.** For the (a) retroelement-free intron clone and (b) *Alu*-free intron clone, graphical representation of the human *TP53* locus is shown on the top: black boxes represent the positions of homology arms used in our UKiS mutagenesis to *TP53*, a horizontal line flanked by two arrowheads represents the target region for the PCR. The orange filled circles denote the position of the heterozygous SNP site, rs12947788. Direct sequencing of the PCR genotyping amplicons indicated double peaks only at rs12947788 on the resultant sequencing chromatograms for all three clones for both mutants.

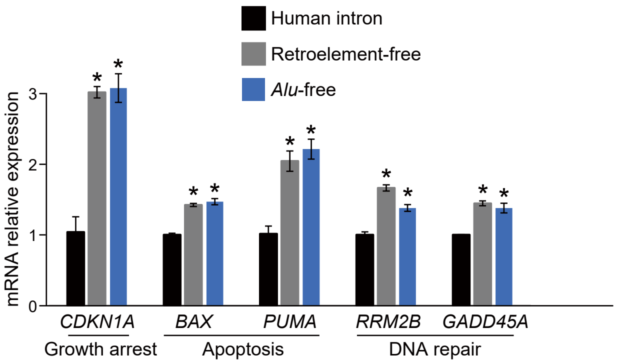


**Supplementary Figure 8 | Evaluating the influence of the retroelements and *Alu* elements in the first intron of *TP53* on the expression of *TP53* downstream genes.**

Real-time RT-PCR for transcriptional expression of five genes that are regulated by *TP53* in the HCT116 cell clones having the *TP53* first intron that are of full-length human, retroelement-free and *Alu*-free. Three different clones of each intron type that were tested in Fig. 6b were used here. As *TP53*-regulating genes, cyclin dependent kinase inhibitor 1A (*CDKN1A* or *p21*), BCL2 associated X (*BAX*), p53 upregulated modulator of apoptosis (*PUMA*), ribonucleotide reductase regulatory TP53 inducible subunit M2B (*RRM2B* or *p53R2*), and growth arrest and DNA damage inducible alpha (*GADD45A*) were examined. 18S rRNA was used as an internal control. Reactions were run in duplicate in three independent experiments. Data represent the mean ± SD, and the p-values were determined by a Welch's t-test. Statistical significance relative to the full-length human intron is denoted with asterisks: $*P < 0.05$

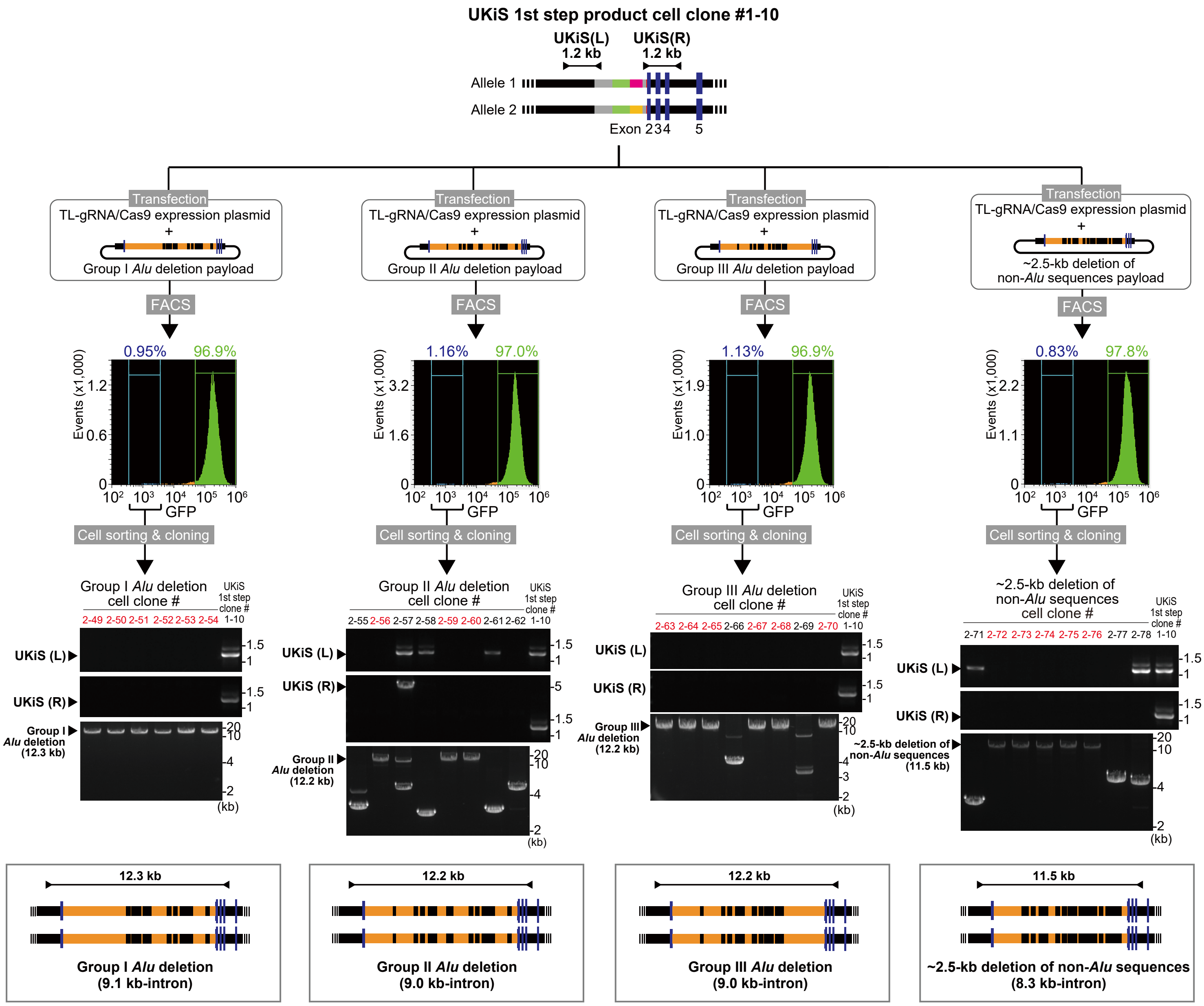


**Supplementary Figure 9 | Replacement of both UKiS donor alleles with the first intron of *TP53* that contains only a subset of *Alu* sequences or that is lacking ~2.5kb of non-*Alu* sequences.** Replacement of both UKiS donor alleles with synthetic introns in which some of the *Alu* sequences or non-*Alu* sequences have been removed. First, the mutating payload plasmids containing the partial deletion of *Alu* sequences or deletion of non-*Alu* sequences from the first intron of *TP53* and the TL-gRNA/Cas9 expression plasmid were transfected into clone #1-10. Thereafter, GFP-negative cells were collected by FACS and cloned. Biallelic substitution of UKiS markers with the mutating payload plasmid was confirmed by junction genotyping PCR, which targeted the regions represented by horizontal lines flanked by arrows in the schematic diagrams of the *TP53* locus after successful replacement. In the agarose gel images, lane numbers of clones that underwent successful recombination in both alleles are in red.

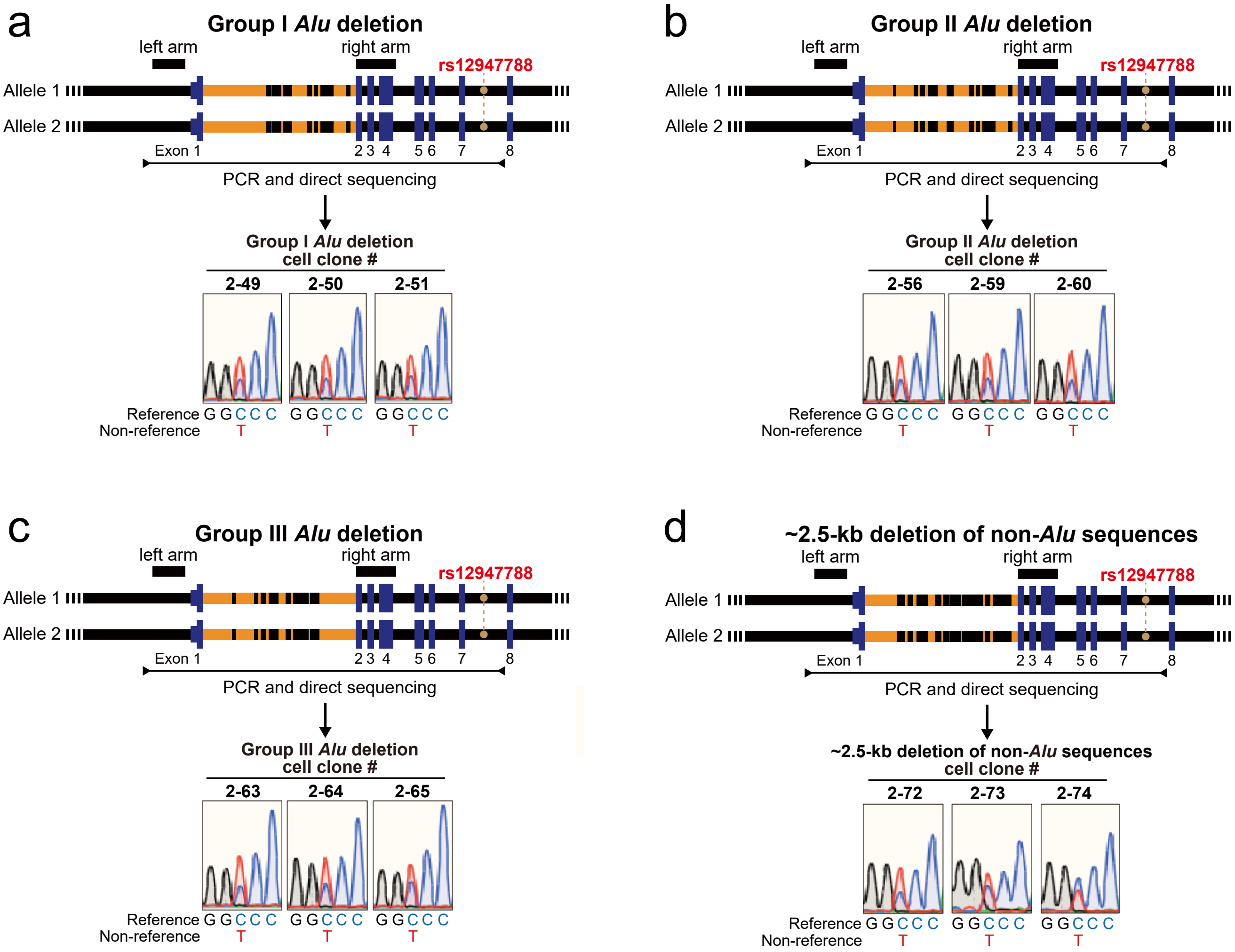


**Supplementary Figure 10 | Validation of biallelic replacement among clones with the first intron of *TP53* that contains only a subset of *Alu* sequences or that is lacking ~2.5kb of non-*Alu* sequences.**

**a, b, c, d.** For each of the (a) Group I *Alu* deletion, (b) Group II *Alu* deletion, and (c) Group III *Alu* deletion clones and the (d) ~2.5-kb deletion of non-*Alu* sequences clones, graphical representation of the human *TP53* gene locus is shown on the top: black boxes represent the positions of homology arms used in our UKiS mutagenesis to *TP53*, a horizontal line flanked by two arrowheads represents the target region for the PCR, and the orange filled circles denote the position of the heterozygous SNP site, rs12947788. Direct sequencing of the PCR genotyping amplicons indicated double peaks only at rs12947788 on the resultant sequencing chromatograms for all three clones of each mutant.

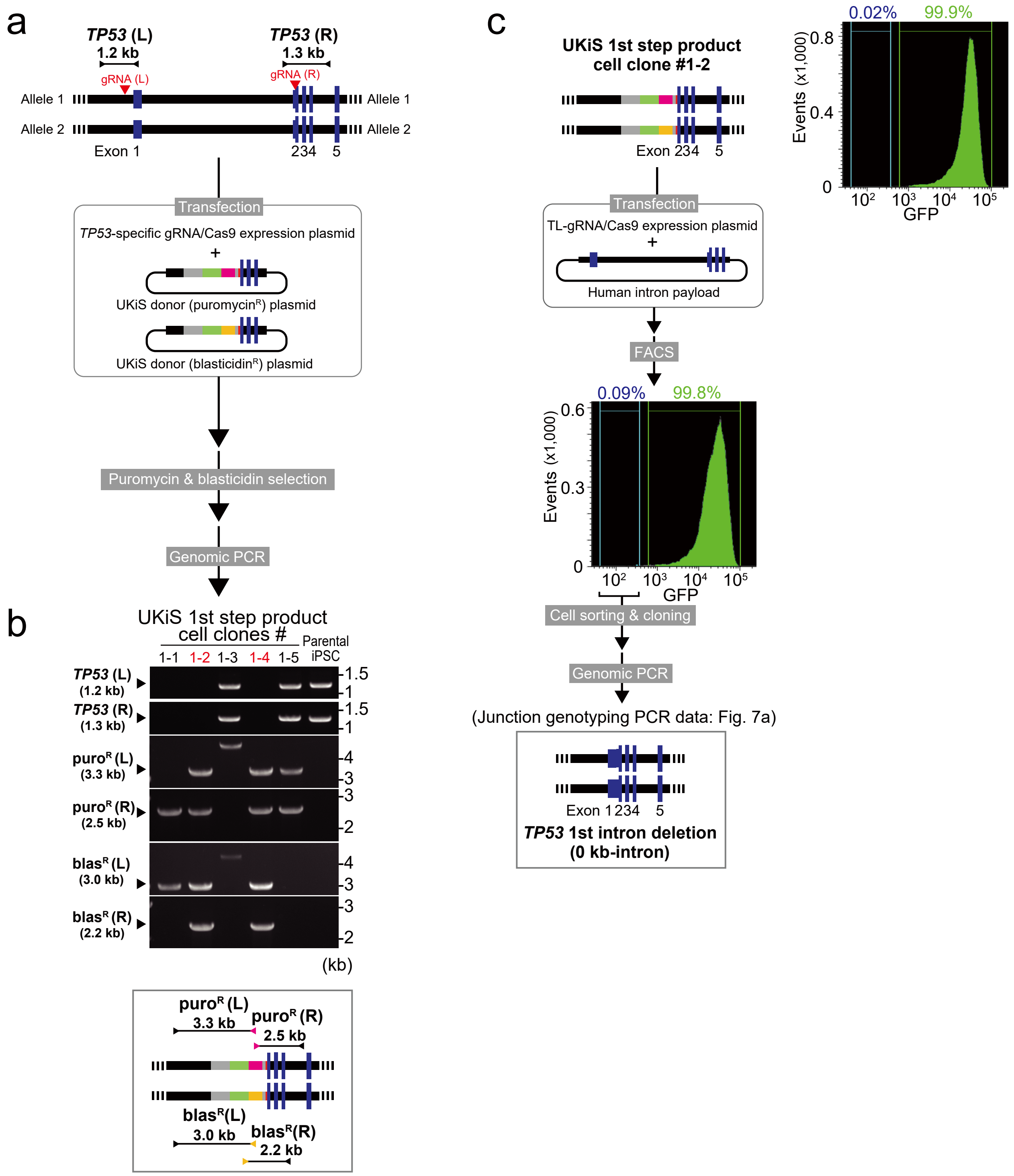


**Supplementary Figure 11 | UKiS deletion of the first intron of *TP53* in iPS cells.**

**a.** Schematic diagram of the first step of UKiS for the first intron of *TP53*. Both UKiS donor plasmids were transfected into iPS cells, leading to isolation of cell clones that had undergone homologous recombination within the *TP53* locus after dual selection with puromycin and blasticidin. Horizontal lines flanked by two arrowheads represent the target regions for the junction genotyping PCR, with the expected length of PCR genotyping amplicons indicated.

**b.** Representative gel images of the junction genotyping PCR to confirm deletion of the first intron of *TP53* and insertion of UKiS markers. Of the five selected clones, two (shown in red) had successful replacement of the intron with the UKiS donor in both alleles.

**c.** Schematic diagram of the second step of UKiS for replacement of both UKiS donor alleles with the *TP53* sequence with the first intron deleted. The payload plasmid with the first intron deleted and the TL-gRNA/Cas9 expression plasmid were transfected into the iPS cell clone #1-2, which was obtained in (b) and mostly GFP-positive as shown by FACS analysis. Thereafter, GFP-negative cells were collected by FACS and cloned. Biallelic substitution of UKiS markers with the mutating payload plasmid was confirmed by junction genotyping PCR. A representative image of the resulting agarose gel electrophoresis of junction PCR amplicons is shown in Figure 7a.

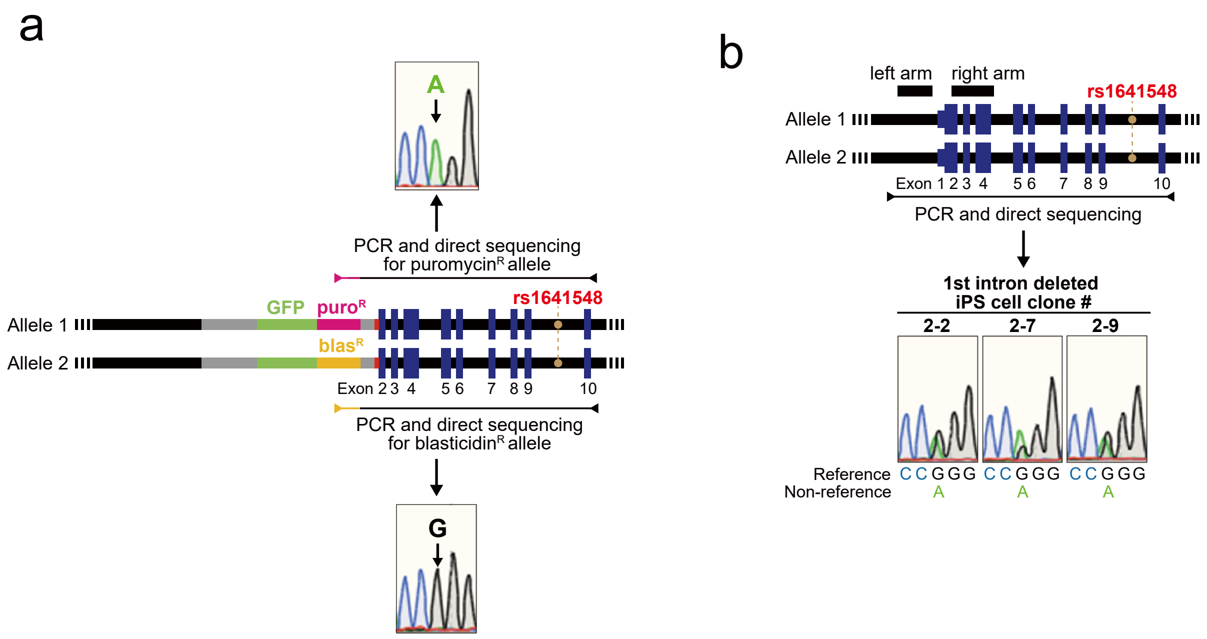


**Supplementary Figure 12 | Validation of biallelic replacement in the second step of UKiS to delete the first intron of *TP53* in iPS cell clones.**

**a.** Genotyping of the iPS cell clone #1-2, which was obtained after the first step of UKiS and was used to create the deletion of the first intron of *TP53* in iPS cell clones. Allele-specific PCR was performed by using primers for puromycin or blasticidin marker sequences, demonstrating that the puromycin and blasticidin alleles had A and G at rs1641548, respectively. Horizontal lines flanked by two arrowheads represent the target region for the PCR of each allele.

**b.** Graphical representation of the human *TP53* locus is shown on the top: black boxes represent the positions of homology arms used in our UKiS mutagenesis to *TP53*, the horizontal line flanked by two arrowheads represents the target region for PCR, and the orange filled circles denote the position of the heterozygous SNP site, rs1641548. Direct sequencing of the PCR genotyping amplicons indicated double peaks only at rs1641548 on the resultant sequencing chromatograms for all three clones of the mutant.

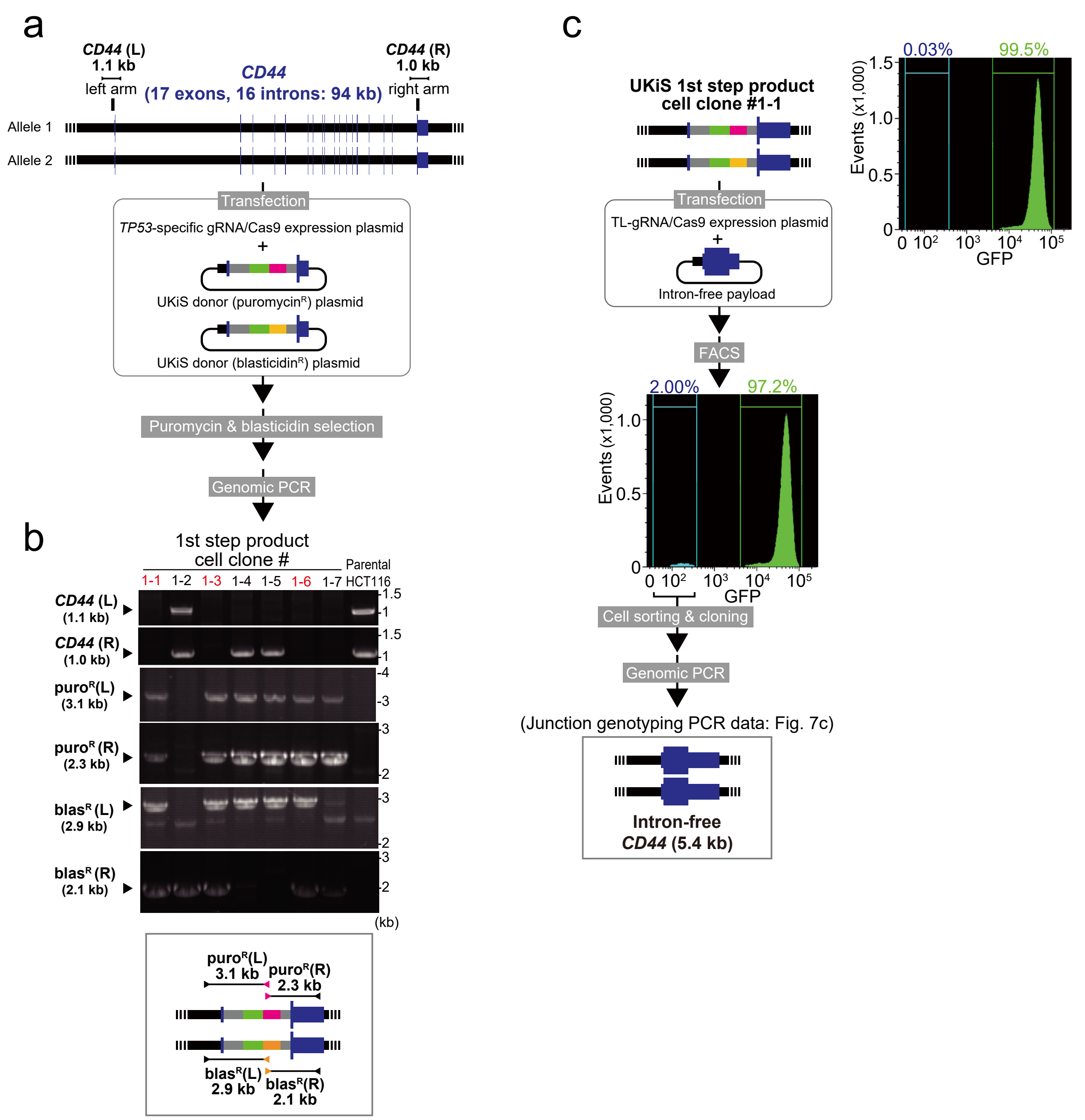


**Supplementary Figure 13 | Deletion of all introns from *CD44* in HCT116 cells by UKiS.**
**a.** Schematic diagram of the first step of UKiS used to generate intron-free *CD44* cell clones. Both UKiS donor plasmids were transfected into HCT116 cells, leading to isolation of cell clones that had undergone homologous recombination within the *CD44* locus after dual selection with puromycin and blasticidin. Horizontal lines flanked by two arrowheads represent the target regions for junction genotyping PCR, with the expected length of PCR genotyping amplicons indicated.
**b.** Representative gel image of the junction genotyping PCR to confirm deletion of *CD44* and insertion of the UKiS markers. Of the seven selected clones, three (shown in red) had successful replacement of the *CD44* sequence with the UKiS donor in both alleles.
**c.** Schematic diagram of the second step of UKiS during which both UKiS donor alleles were replaced with the intron-free *CD44* sequence. First, the intron-free *CD44* payload plasmid and the TL-gRNA/Cas9 expression plasmid were transfected into clone #1-1, which was obtained in (b) and mostly GFP-positive as shown by FACS analysis. Thereafter, GFP-negative cells were collected by FACS and cloned. Biallelic substitution of UKiS markers with the mutating payload plasmid was confirmed by junction genotyping PCR. A representative image of the resulting agarose gel electrophoresis of junction PCR amplicons is shown in Figure 7c.

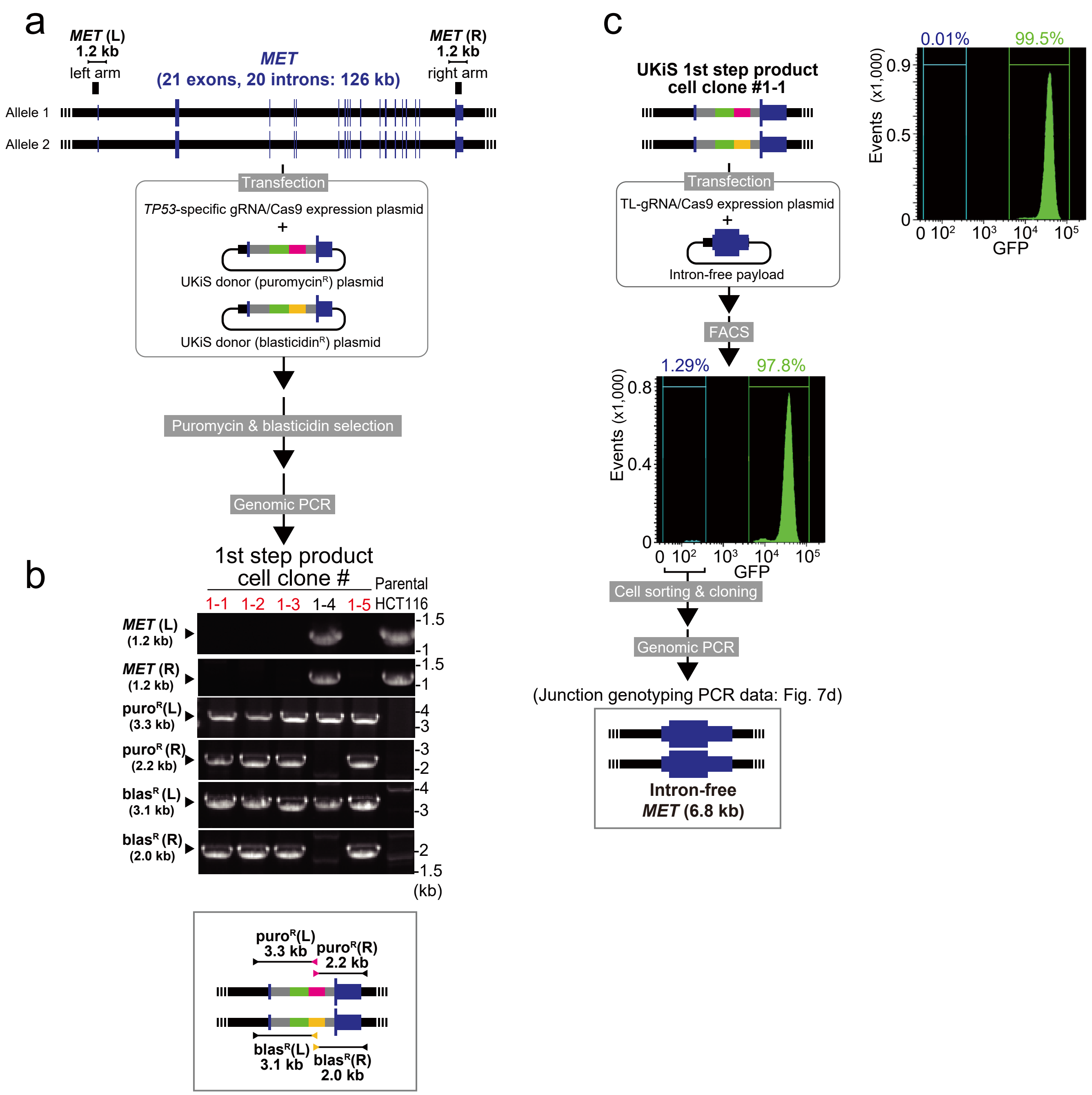


**Supplementary Figure 14 | Deletion of all introns from *MET* in HCT116 cells by UKiS.**

**a.** Schematic diagram of the first step of UKiS used to generate intron-free *MET* cell clones. Both UKiS donor plasmids were transfected into HCT116 cells, leading to isolation of cell clones that had undergone homologous recombination within the *MET* locus after dual selection with puromycin and blasticidin. Horizontal lines flanked by two arrowheads represent the target regions for junction genotyping PCR, with the expected length of PCR genotyping amplicons indicated.

**b.** Representative gel image of the junction genotyping PCR to confirm deletion of *MET* and insertion of the UKiS markers. Of the five selected clones, two (shown in red) had successful replacement of the *MET* sequence with the UKiS donor in both alleles.

**c.** Schematic diagram of the second step of UKiS during which both UKiS donor alleles were replaced with the intron-free *MET* sequence. First, the intron-free *MET* payload plasmid and the TL-gRNA/Cas9 expression plasmid were transfected into clone #1-1, which was obtained in (b) and mostly GFP-positive as shown by FACS analysis. Thereafter, GFP-negative cells were collected by FACS and cloned. Biallelic substitution of UKiS markers with the mutating payload plasmid was confirmed by junction genotyping PCR. A representative image of the resulting agarose gel electrophoresis of junction PCR amplicons is shown in Figure 7d.

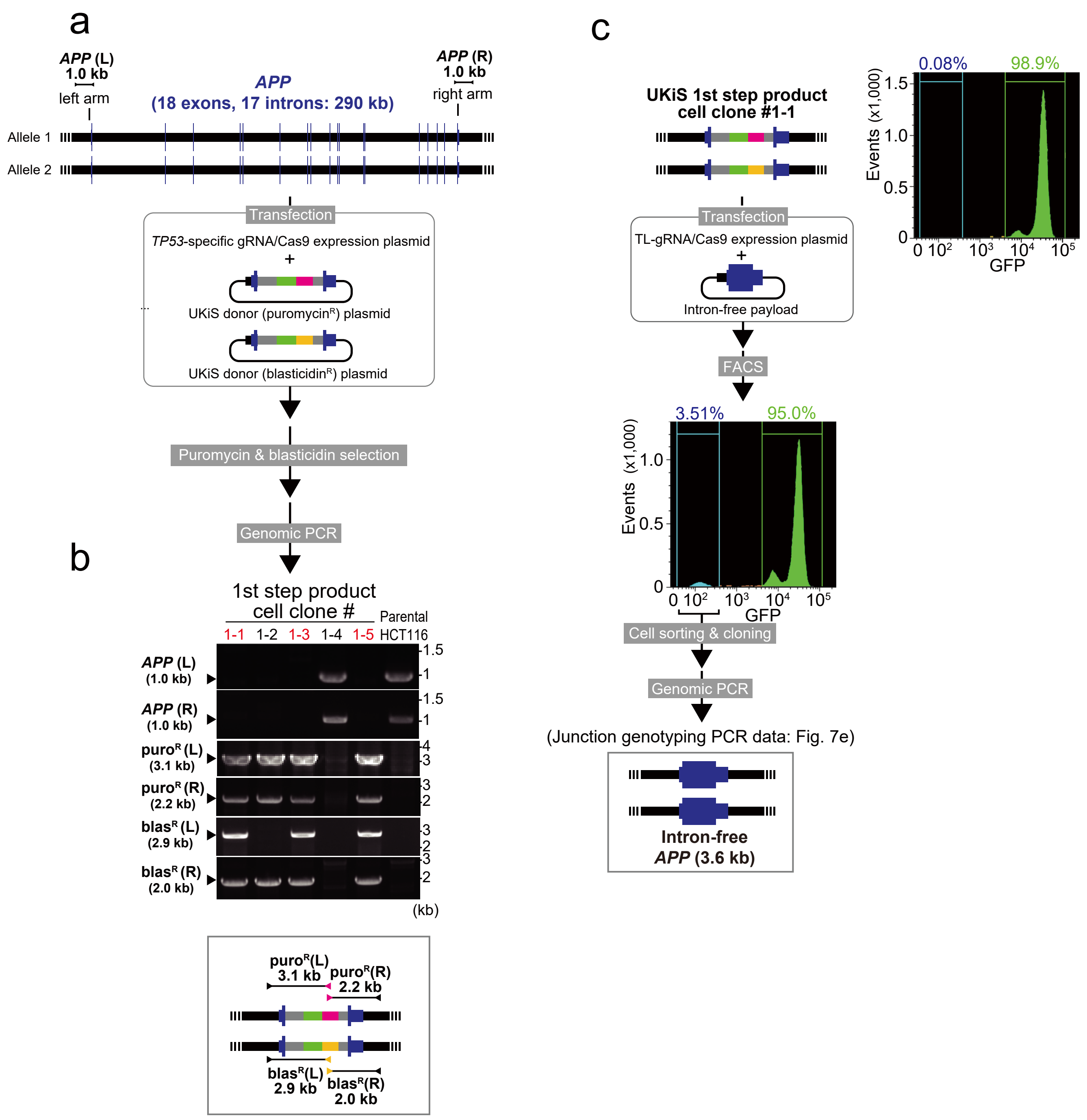


**Supplementary Figure 15 | Deletion of all introns from *APP* in HCT116 cells by UKiS.**

**a.** Schematic diagram of the first step of UKiS used to generate intron-free *APP* cell clones. Both UKiS donor plasmids were transfected into HCT116 cells, leading to isolation of cell clones that had undergone homologous recombination within the *APP* locus after dual selection with puromycin and blasticidin. Horizontal lines flanked by two arrowheads represent the target regions for junction genotyping PCR, with the expected length of PCR genotyping amplicons indicated.

**b.** Representative gel image of the junction genotyping PCR to confirm deletion of *APP* and insertion of the UKiS markers. Of the five selected clones, three (shown in red) had successful replacement of the *APP* sequence with the UKiS donor in both alleles.

**c.** Schematic diagram of the second step of UKiS during which both UKiS donor alleles were replaced with the intron-free *APP* sequence. First, the intron-free *APP* payload plasmid and the TL-gRNA/Cas9 expression plasmid were transfected into clone #1-1, which was obtained in (b) and mostly GFP-positive as shown by FACS analysis. Thereafter, GFP-negative cells were collected by FACS and cloned. Biallelic substitution of UKiS markers with the mutating payload plasmid was confirmed by junction genotyping PCR. A representative image of the resulting agarose gel electrophoresis of junction PCR amplicons is shown in Figure 7e.

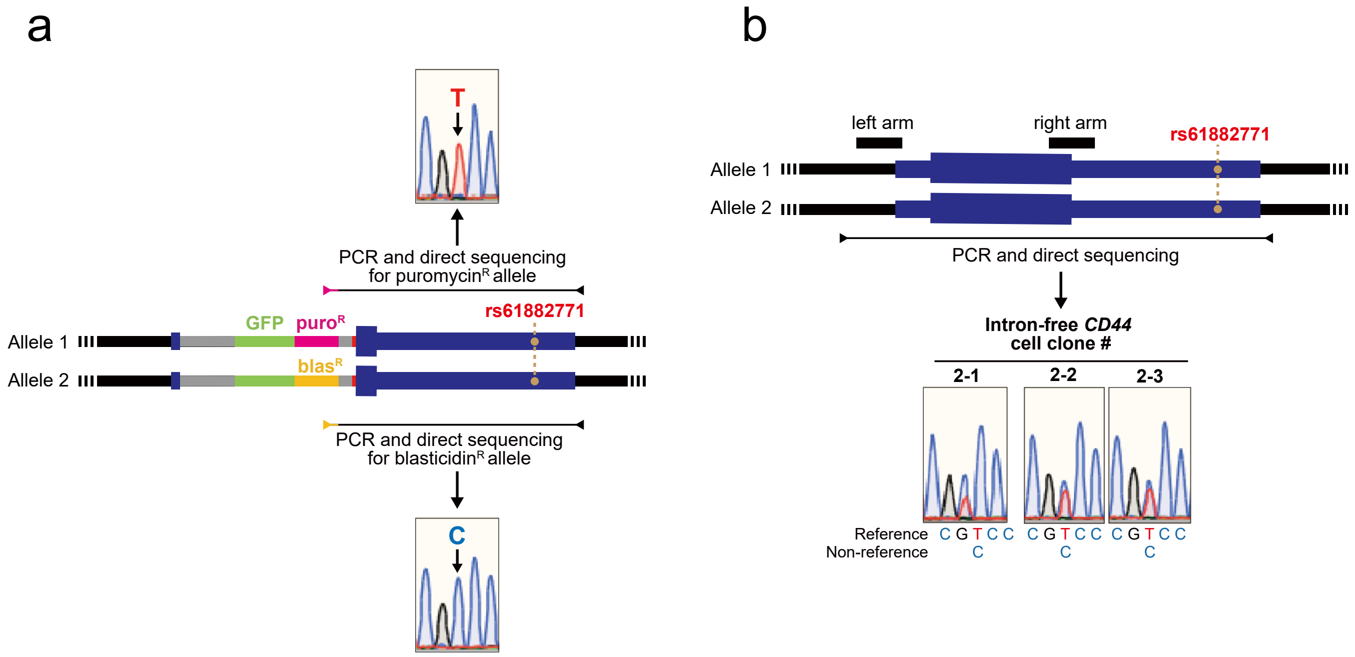


**Supplementary Figure 16 | Validation of biallelic replacement of the *CD44* locus in the first and second step of UKiS to generate intron-free *CD44* cell clones.**

**a.** Genotyping of clone #1-1 in the *CD44* region after the first step of UKiS. Clone #1-1 was then used to create the intron-free *CD44* cell clones. Allele-specific PCR was performed by using primers for puromycin or blasticidin marker sequences, demonstrating that the puromycin and blasticidin alleles have a T and C at rs61882771, respectively. Horizontal lines flanked by two arrowheads represent the target region for the PCR of each allele.

**b.** Graphical representation of the intron-free *CD44* locus is shown on the top: black boxes represent the positions of homology arms used in our UKiS mutagenesis to *CD44*, the horizontal line flanked by two arrowheads represents the target region for PCR, and the orange filled circles denote the position of the heterozygous SNP site, rs61882771. Direct sequencing of the PCR genotyping amplicons indicated double peaks only at rs61882771 on the resultant sequencing chromatograms for all three clones of this mutant.

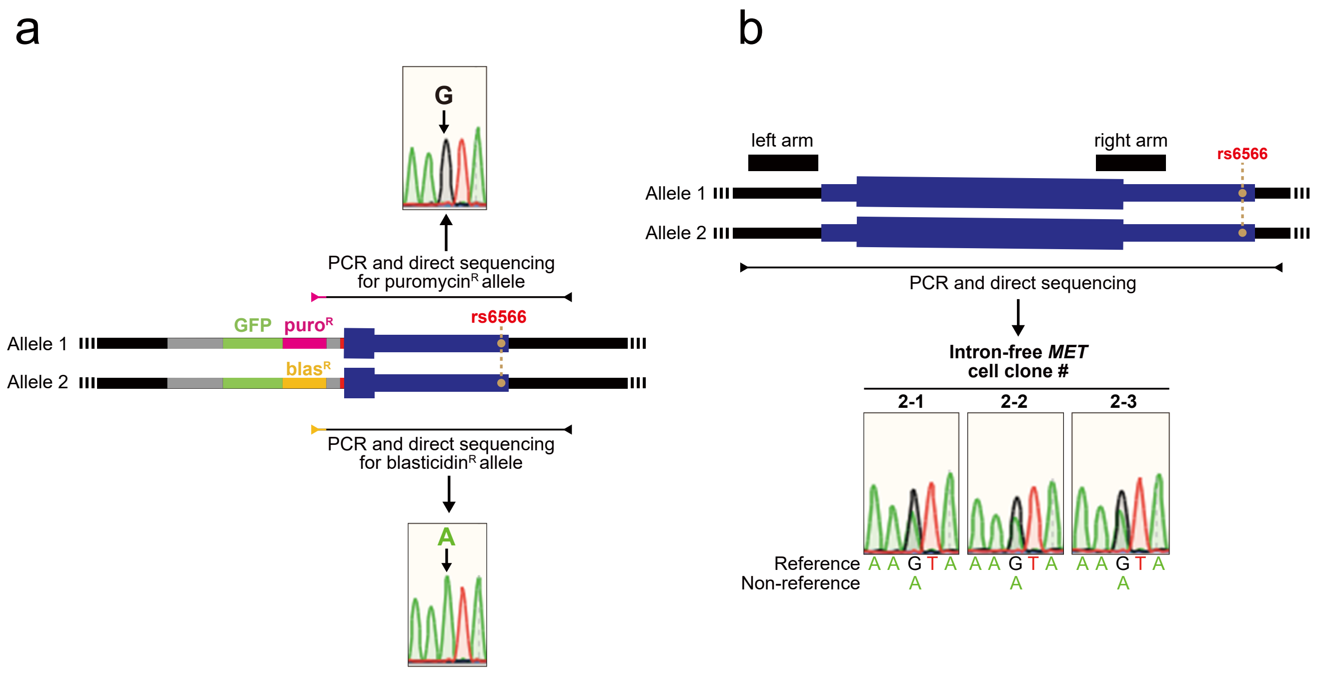


**Supplementary Figure 17 | Validation of biallelic replacement of the *MET* locus in the first and second step of UKiS to generate intron-free *MET* cell clones.**

**a.** Genotyping of clone #1-1 in the *MET* region after the first step of UKiS. Clone #1-1 was then used to create the intron-free *MET* cell clones. Allele-specific PCR was performed by using primers for puromycin or blasticidin marker sequences, demonstrating that the puromycin and blasticidin alleles have a G and A at rs6566, respectively. Horizontal lines flanked by two arrowheads represent the target region for the PCR of each allele.

**b.** Graphical representation of the intron-free *MET* locus is shown on the top: black boxes represent the positions of homology arms used in our UKiS mutagenesis to *MET*, the horizontal line flanked by two arrowheads represents the target region for PCR, and the orange filled circles denote the position of the heterozygous SNP site, rs6566. Direct sequencing of the PCR genotyping amplicons indicated double peaks only at rs6566 on the resultant sequencing chromatograms for all three clones of this mutant.

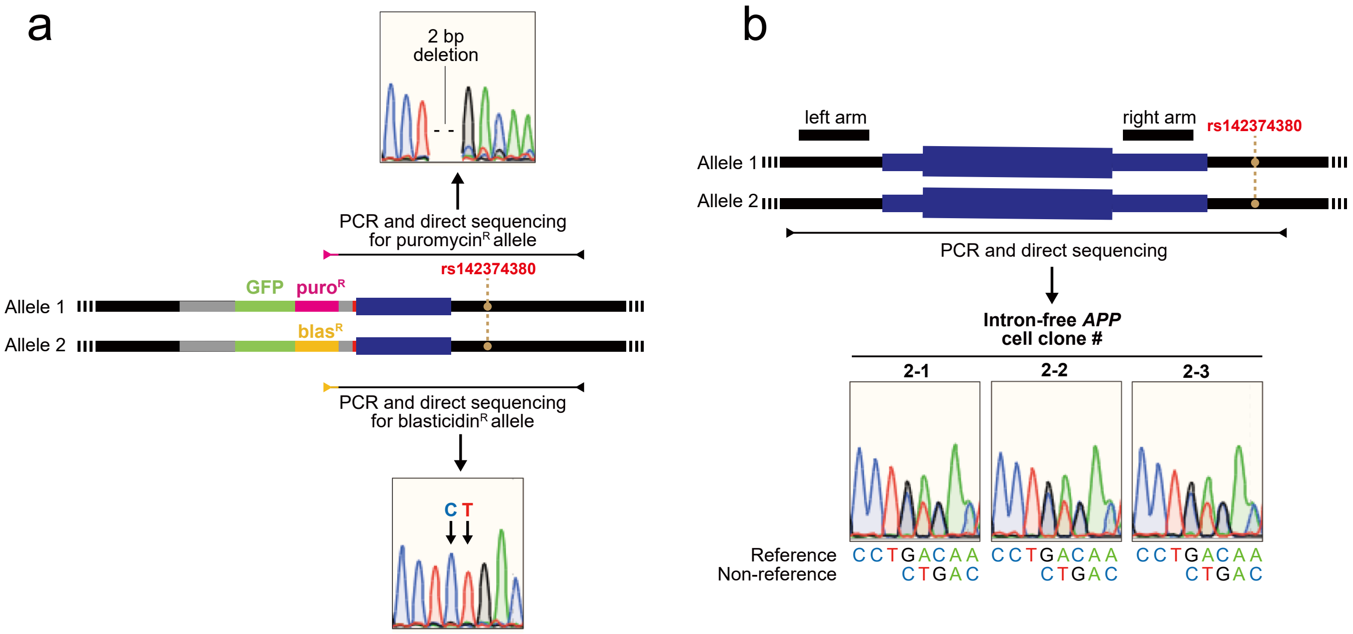


**Supplementary Figure 18 | Validation of biallelic replacement of the *APP* locus in the first and second step of UKiS to generate intron-free *APP* cell clones.**

**a.** Genotyping of clone #1-1 in the *APP* region after the first step of UKiS. Clone #1-1 was then used to create the intron-free *APP* cell clones. Allele-specific PCR was performed by using primers for puromycin or blasticidin marker sequences, demonstrating that the puromycin and blasticidin alleles have a 2-bp deletion and a CT at rs142374380, respectively. Horizontal lines flanked by two arrowheads represent the target region for the PCR of each allele.

**b.** Graphical representation of the intron-free *APP* gene locus is shown on the top: black boxes represent the positions of homology arms used in our UKiS mutagenesis to *APP*, the horizontal line flanked by two arrowheads represents the target region for PCR, and the orange filled circles denote the position of the heterozygous SNP site, rs142374380. Direct sequencing of the PCR genotyping amplicons indicated an equivalent mixture of the 2-bp deletion and CT peaks at rs142374380 on the resultant sequencing chromatograms for all three clones of this mutant.

### Supplementary Table 1. Primers used for genotyping junction PCR

| Fig. # | Target | Forward primer | Reverse primer |
|---|---|---|---|
| Fig. 2c<br>Supplementary Figs. 2b and 11b | *TP53* (L) | TGCCCCGTTGTTATCCTTAC | GATGCAGAAGAGGTGCAAGA |
| | *TP53* (R) | GAGCCGCAGTCAGATCCTAG | GAAGACGGCAGCAAAGAAAC |
| | puro$^R$ (L) (*TP53*) | TGCCCCGTTGTTATCCTTAC | GCTCGTAGAAGGGGAGGTTG |
| | puro$^R$ (R) (*TP53*) | GTCACCGAGCTGCAAGAAC | GAAGACGGCAGCAAAGAAAC |
| | blas$^R$ (L) (*TP53*) | TGCCCCGTTGTTATCCTTAC | GCTTCAATATGTACTGCCGAAA |
| | blas$^R$ (R) (*TP53*) | GAAGCCATTGCGATTGGTAG | GAAGACGGCAGCAAAGAAAC |
| Figs. 3b-e and 7a<br>Supplementary Figs. 6 and 9 | eintire *TP53* first intron | TGCCCCGTTGTTATCCTTAC | GAAGACGGCAGCAAAGAAAC |
| | UKiS(L) (*TP53*) | TGCCCCGTTGTTATCCTTAC | GGGCTGTCCCTCATAAAAGT |
| | UKiS(R) (*TP53*) | TCACGTAAGTAGAACATGAAATAACC | GAAGACGGCAGCAAAGAAAC |
| Fig. 7c | entire *CD44* | AGTGGATGGACAGGAGGATG | GAGTGGGTCTGAGTGGGAAC |
| Fig. 7d | entire *MET* | TGAAATCACTCTTATGTAACCTCTGG | TGCAGGTATAGGCAGTGACAAG |
| Fig. 7e | entire *APP* | GGGGAGCTGGTACAGAAATG | CTCTTCTCCCCACCCAAAA |
| Supplementary Fig. 13b | *CD44* (L) | AGTGGATGGACAGGAGGATG | ACCCATCTTGCTGCCCAGG |
| | *CD44* (R) | ATGGAGCTGTGGAGGACAGA | GAGTGGGTCTGAGTGGGAAC |
| | puro$^R$ (L) (*CD44*) | AGTGGATGGACAGGAGGATG | GCTCGTAGAAGGGGAGGTTG |
| | puro$^R$ (R) (*CD44*) | GTCACCGAGCTGCAAGAAC | GAGTGGGTCTGAGTGGGAAC |
| | blas$^R$ (L) (*CD44*) | AGTGGATGGACAGGAGGATG | GCTTCAATATGTACTGCCGAAA |
| | blas$^R$ (R) (*CD44*) | GAAGCCATTGCGATTGGTAG | GAGTGGGTCTGAGTGGGAAC |
| Supplementary Fig. 14b | *MET* (L) | TGAAATCACTCTTATGTAACCTCTGG | CCGGGCATCGGCGCGCGCGGCCC |
| | *MET* (R) | ACCCACTGTTTGAGAATGATG | TGCAGGTATAGGCAGTGACAAG |
| | puro$^R$ (L) (*MET*) | TGAAATCACTCTTATGTAACCTCTGG | GCTCGTAGAAGGGGAGGTTG |
| | puro$^R$ (R) (*MET*) | GTCACCGAGCTGCAAGAAC | TGCAGGTATAGGCAGTGACAAG |
| | blas$^R$ (L) (*MET*) | TGAAATCACTCTTATGTAACCTCTGG | GCTTCAATATGTACTGCCGAAA |
| | blas$^R$ (R) (*MET*) | GAAGCCATTGCGATTGGTAG | TGCAGGTATAGGCAGTGACAAG |
| Supplementary Fig. 15b | *APP* (L) | GGGGAGCTGGTACAGAAATG | CAGGATCAGGGAAAGGTGAG |
| | *APP* (R) | GAACGGCTACGAAAATCCAA | CTCTTCTCCCCACCCAAAA |
| | puro$^R$ (L) (*APP*) | GGGGAGCTGGTACAGAAATG | GCTCGTAGAAGGGGAGGTTG |
| | puro$^R$ (R) (*APP*) | GTCACCGAGCTGCAAGAAC | CTCTTCTCCCCACCCAAAA |
| | blas$^R$ (L) (*APP*) | GGGGAGCTGGTACAGAAATG | GCTTCAATATGTACTGCCGAAA |
| | blas$^R$ (R) (*APP*) | GAAGCCATTGCGATTGGTAG | CTCTTCTCCCCACCCAAAA |

### Supplementary Table 2. Primers used for SNP typing

#### Primers used for PCR

| Fig. # | Target | Forward primer | Reverse primer |
|---|---|---|---|
| Fig. 4a | From *TP53* 4th intron to 7th intron | GGCTTTTATCCATCCCATCA | AGCTGTTCCGTCCCAGTAGA |
| Fig. 4b | From puromycinR sequence to *TP53* 7th intron | GTCACCGAGCTGCAAGAAC | AGCTGTTCCGTCCCAGTAGA |
| | From blasticidinR sequence to *TP53* 7th intron | GAAGCCATTGCGATTGGTAG | AGCTGTTCCGTCCCAGTAGA |
| Fig. 4c<br>Supplementary Figs. 5, 7 and 10 | From upstream of *TP53* 1st exon to 7th intron | TGCCCCGTTGTTATCCTTAC | AGCTGTTCCGTCCCAGTAGA |
| Supplementary Fig. 12a | From puromycin resistance gene sequence to *TP53* 11th intron | GTCACCGAGCTGCAAGAAC | AGCTGCCTTTGACCATGAAG |
| | From blasticidin resistance gene sequence to *TP53* 11th intron | GAAGCCATTGCGATTGGTAG | AGCTGCCTTTGACCATGAAG |
| Supplementary Fig. 12b | From upstream of *TP53* 1st exon to 10th intron | TGCCCCGTTGTTATCCTTAC | AGCTGCCTTTGACCATGAAG |
| Supplementary Fig. 16a | From puromycin resistance gene sequence to downstream of *CD44* last exon | GTCACCGAGCTGCAAGAAC | AAGTTGATGGAAACCCATTTTG |
| | From blasticidin resistance gene sequence to downstream of *CD44* last exon | GAAGCCATTGCGATTGGTAG | AAGTTGATGGAAACCCATTTTG |
| Supplementary Fig. 16b | From upstream of *CD44* 1st exon to downstream of last exon | AGTGGATGGACAGGAGGATG | AAGTTGATGGAAACCCATTTTG |
| Supplementary Fig. 17a | From puromycin resistance gene sequence to downstream of *MET* last exon | GTCACCGAGCTGCAAGAAC | CCAAACCCTTGGAAAGACAG |
| | From blasticidin resistance gene sequence to downstream of *MET* last exon | GAAGCCATTGCGATTGGTAG | CCAAACCCTTGGAAAGACAG |
| Supplementary Fig. 17b | From upstream of *MET* 1st exon to downstream of last exon | TGAAATCACTCTTATGTAACCTCTGG | CCAAACCCTTGGAAAGACAG |
| Supplementary Fig. 18a | From puromycin resistance gene sequence to downstream of *APP* last exon | GTCACCGAGCTGCAAGAAC | ATTTGGCAGCACTTCAGTTT |
| | From blasticidin resistance gene sequence to downstream of APP last exon | GAAGCCATTGCGATTGGTAG | ATTTGGCAGCACTTCAGTTT |
| Supplementary Fig. 18b | From upstream of *APP* 1st exon to downstream of last exon | GGGGAGCTGGTACAGAAATG | ATTTGGCAGCACTTCAGTTT |

#### Primers used for direct sequencing

| Fig. # | Target | Primer |
|---|---|---|
| Fig. 4a | Heterozygous SNP candidates in *TP53* locus | GGGATTCCTCTTCACCCTTT |
| | Heterozygous SNP candidates in *TP53* locus | AGGTGTAGACGCCAACTCTCTC |
| | Heterozygous SNP candidates in *TP53* locus | GTGTGGTGGTGCCCTATGA |
| | Heterozygous SNP candidates in *TP53* locus | ATCCTGGCTAACGGTGAAAC |
| | Heterozygous SNP candidates in *TP53* locus | CCTCACCATCATCACACTGG |
| | Heterozygous SNP candidates in *TP53* locus | GGCGCACAGAGGAAGAGA |
| | Heterozygous SNP candidates in *TP53* locus | GCAGTGATGCCTCAAAGACA |
| | Heterozygous SNP candidates in *TP53* locus | CTTGTGATCTGCCTGCCTTG |
| | Heterozygous SNP candidates in *TP53* locus | CAACTACAGGCCTGCACCAC |
| | Heterozygous SNP candidates in *TP53* locus | TGCCCATGCTGGTATCAAAC |
| | Heterozygous SNP candidates in *TP53* locus | TTGCAGGGAGCCAAGATG |
| | Heterozygous SNP candidates in *TP53* locus | AGCTGCCTTTGACCATGAAG |
| Fig. 4b and 4c<br>Supplementary Fig. 5, 7, 10 | rs12947788 | AGCTGTTCCGTCCCAGTAGA |
| Supplementary Fig. 12 | rs1641548 | AGCTGCCTTTGACCATGAAG |
| Supplementary Fig. 16 | rs61882771 | CCATTTTCAGTGGTCTGGATT |
| Supplementary Fig. 17 | rs6566 | TCTGCTCTGTGGAAAGAAAGA |
| Supplementary Fig. 18 | rs142374380 | ATTTGGCAGCACTTCAGTTT |

**Supplementary Table 3. Primers used for RT-PCR**

| Fig. # | Target | Forward primer | Reverse primer |
|---|---|---|---|
| Fig. 5a | *TP53* | CTCAAGACTGGCGCTAAAAG | CTGCCCTGGTAGGTTTTCTG |
| Figs. 5a and 7f-h | 18S rRNA | ATTAATCAAGAACGAAAGTCGGAGGT | TTTAAGTTTCAGCTTTGCAACCATACT |
| Fig. 7f | *CD44* | CATCCTCGTCCCGTCCTC | CCATGAGATTTGGCTGAGTG |
| Fig. 7g | *MET* | GAGCGCCTCAGTCTGGTC | TGCAGGTATAGGCAGTGACAAG |
| Fig. 7h | *APP* | GATCCCACTCGCACAGCA | TTCTCCCCACCCAAAATTAC |

### Supplementary Table 4. All primers and oligonucleotides used for plasmid construction and sequencing, and the constructed plasmid

#### Primers and oligonucleotides used for plasmid construction, and constructed plasmid

| Plasmid | Description | Backbone | PCR target | PCR template | Forward primer | Reverse primer |
|---|---|---|---|---|---|---|
| HR110PA-1_GFP | HR110PA-1 plasmid (RFP coding sequence of HR110PA-1 was replaced by GFP) | HR110PA-1 | Left insulator, EF1 promoter and part of backbone | HR110PA-1 | GGTGATGACGGTGAAAACCT | CCCTTGCTCACGGTACCCATGGTGGCGATATCGTAGGCG |
| | | | T2A, Puromycin resistance gene, right insulator and part of backbone | HR110PA-1 | GAGGGCAGAGGAAGTCTTCT | AGGTTTTCACCGTCATCACC |
| | | | GFP | HR110PA-1 | ATGGGTACCGTGAGCAAGG | CTCCACGTCACCGCATGTTAGAAGACTTCCTCTGCCCTCCTTGTACAGCTCGTCCATG |
| HR110PA-1_GFP_blasR | HR110PA-1 plasmid (Puromycin resistance gene of HR110PA-1_GFP was replaced by that of blasticidin resistance gene) | HR110PA-1 | Blasticidin resistance gene | pCISP310B | ATGAAAACATTTAACATTTCTCAAC | TTTTAATTTCGGGTATATTTGAGTGGA |
| | | | Left insulator, EF1 promoter, GFP and part of backbone | HR110PA-1_GFP | GGTGATGACGGTGAAAACCT | AATGTTAAATGTTTTCATAGGGCCGGGATTCTCCTCCA |
| | | | T2A, right insulator and part of | HR110PA-1_GFP | AAATATACCCGAAATTAAAATCAACCTCTGGAC | AGGTTTTCACCGTCATCACC |
| pTO455 | UKiS donor (puromycin resistance gene) for *TP53* locus | HR110PA-1 | *TP53* left arm | HCT116 genome | GTAAAACGACGGCCAGTGAATTCTTCCCCAGAGGTATCTTCCA | AGCGTAGGCGCTTCTCGCCAAGAT |
| | | | *TP53* right arm | HCT116 genome | GCGCAACGCGATCGCGTAAGGGCTGAGTCAGGAAACATTTTCAG | CTTGCATGCAGTCGACGGGGATCCAAGTTCTGCATCCCCAGGAG |
| pTO456 | UKiS donor (blasticidin resistance gene) for *TP53* locus | HR110PA-1 | *TP53* left arm | HCT116 genome | GTAAAACGACGGCCAGTGAATTCTTCCCCAGAGGTATCTTCCA | AGCGTAGGCGCTTCTCGCCAAGAT |
| | | | *TP53* right arm | HCT116 genome | GCGCAACGCGATCGCGTAAGGGCTGAGTCAGGAAACATTTTCAG | CTTGCATGCAGTCGACGGGGATCCAAGTTCTGCATCCCCAGGAG |
| pTO467 | Mutating payload for creating human *TP53* 1st intron restored cell | HR110PA-1 | *TP53* left arm to 1st exon | HCT116 genome | GTAAAACGACGGCCAGTGAATTCTTCCCCAGAGGTATCTTCCA | CCAATCCAGGGAAGCGTGT |
| | | | *TP53* 2nd exon to right arm | HCT116 genome | CAGCCAGACTGCCTTCCGG | CTTGCATGCAGTCGACGGGGATCCAAGTTCTGCATCCCCAGGAG |
| | | | Human *TP53* 1st intron | HCT116 genome | GTAAAACGACGGCCAGTGAATTCTTCCCCAGAGGTATCTTCCA | CTTGCATGCAGTCGACGGGGATCCAAGTTCTGCATCCCCAGGAG |
| pTO469 | Mutating payload for creating mouse *Tp53* 1st intron replaced cell | HR110PA-1 | *TP53* left arm to 1st exon | HCT116 genome | GTAAAACGACGGCCAGTGAATTCTTCCCCAGAGGTATCTTCCA | CCAATCCAGGGAAGCGTGT |
| | | | *TP53* 2nd exon to right arm | HCT116 genome | CAGCCAGACTGCCTTCCGG | CTTGCATGCAGTCGACGGGGATCCAAGTTCTGCATCCCCAGGAG |
| | | | Mouse *Tp53* 1st intron | Mouse genome | ACACGCTTCCCTGGATTGGGTAAGTAATTGATGAGCGTGA | CGGAAGGCAGTCTGGCTGCTGTAGAGAGAAGAGATTGTG |
| pTO470 | Mutating payload for creating zebrafish *tp5*3 1st intron replaced cell | HR110PA-1 | *TP53* left arm to 1st exon | HCT116 genome | GTAAAACGACGGCCAGTGAATTCTTCCCCAGAGGTATCTTCCA | CCAATCCAGGGAAGCGTGT |
| | | | *TP53* 2nd exon to right arm | HCT116 genome | CAGCCAGACTGCCTTCCGG | CTTGCATGCAGTCGACGGGGATCCAAGTTCTGCATCCCCAGGAG |
| | | | Zebrafish *tp53* 1st intron | Zebrafish genome | ACACGCTTCCCTGGATTGGGTAACAAAGCGAAACTTTTATTG | CCGGAAGGCAGTCTGGCTGCTGTGAAATTATAAACACGAAAG |
| pTO471 | Mutating payload for creating *TP53* 1st intron deleted cell | HR110PA-1 | *TP53* left arm to 1st exon | HCT116 genome | GTAAAACGACGGCCAGTGAATTCTTCCCCAGAGGTATCTTCCA | CCAATCCAGGGAAGCGTGT |
| | | | *TP53* 2nd exon to right arm | HCT116 genome | CAGCCAGACTGCCTTCCGG | CTTGCATGCAGTCGACGGGGATCCAAGTTCTGCATCCCCAGGAG |
| | | | From *TP53* 1st exon to 2nd exon excluding 1st intron | HCT116 cDNA | GATGGGATTGGGGTTTTCCC | CCAATCCAGGGAAGCGTGTC |

**(Continued)**

| Plasmid | Description | Backbone | PCR target | PCR template | Forward primer | Reverse primer |
|---|---|---|---|---|---|---|
| pTO468 | Mutating payload for creating retroelement-free *TP53* 1st intron cell | HR110PA-1 | Retroelement-free fragment #1 including left arm | pTO467 | GTAAAACGACGGCCAGTGAATTCTTCCCCAGAGGTATCTTCCA | TTTAGGAAAAAGATGACGTAAGTACG |
| | | | Retroelement-free fragment #2 | | CGTACTTACGTCATCTTTTTCCTAAACACTGGACATATAGGCCTTC | CTAGTACTCTGTGTATTATGGGAATTCTGCAATTGTTCTATTTCACTTG |
| | | | Retroelement-free fragment #3 | | CCCATAATACACAGAGTACTAGGTAAAAGAAAAGAGCCTGCCATT | CTACTGTAAGTGTTTGTTACACCACTTTTCTGTTGTTTGCACTG |
| | | | Retroelement-free fragment #4 | | GTGCAAACAACAGAAAAGTGGTGTAACAAACACTTACAGTAG | CAAAGGCTTTGCCATGTTTCGCCATGAGGATGCTCTCTTTCTTTTTC |
| | | | Retroelement-free fragment #5 | | GAAAGAGAGCATCCTCATGGCGAAACATGGCAAAGCCTTTGAAAG | GTCATCCTATTTTAATTCACATCAGAATAACACACAAGCCTGTTATA |
| | | | Retroelement-free fragment #6 | | AACAGGCTTGTGTGTTATTCTGATGTGAATTAAAATAGGATGAC | TCCTGTGGAGCAGGAAAAGATCCCACCTCCTGTTAACAAGG |
| | | | Retroelement-free fragment #7 including right arm | | CCTTGTTAACAGGAGGTGGGATCTTTTCCTGCTCCACAGGA | CTTGCATGCAGTCGACGGGGATCCAAGTTCTGCATCCCCAGGAG |
| pTO504 | Mutating payload for creating *Alu*-free *TP53* 1st intron cell | HR110PA-1 | *Alu*-free fragment #1 including left HA arm | pTO467 | GTAAAACGACGGCCAGTGAATTCTTCCCCAGAGGTATCTTCCA | CACATAATGTATGATTCCCCCCTGATTCCATTCTATATGAAGTTCTCC |
| | | | *Alu*-free fragment #2 | | GGAGAACTTCATATAGAATGGAATCAGGGGGGAATCATACATTATGTG | GGAGATAAATAGAAAATAGCACTAAGATAGCACTAAAATGGTAGCCCTA |
| | | | *Alu*-free fragment #3 | | TAGGGCTACCATTTTAGTGCTATCTTAGTGCTATTTTCTATTTATCTCC | AACAAGTTGGGAAATATTCCTCTATGTTCCTCTGTTTTCTGAAGAG |
| | | | *Alu*-free fragment #4 | | CTCTTCAGAAAACAGAGGAACATAGAGGAATATTTCCCAACTTGTT | TAAACATTATTTGGTTAGAATGTATATTTATCCTTTTTTCTATATTACC |
| | | | *Alu*-free fragment #5 | | GGTAATATAGAAAAAAGGATAAATATACATTCTAACCAAATAATGTTTA | TCTTAGAGGAAAAAGCGTTGAGTTAGCACTGAGTTTTCTACCATTA |
| | | | *Alu*-free fragment #6 | | TAATGGTAGAAAACTCAGTGCTAACTCAACGCTTTTTCCTCTAAGA | TATTTCTTCCTTTCTAATCGTTATTACTAATCACTCATTATTTCTTTTTCTT |
| | | | *Alu*-free fragment #7 | | CAGGCAAGAAAAAGAAATAATGAGTGATTAGTAATAACGATTAGAAAGGA | TTTTTGAGATAGAGTTTCACCCATTAGGACATGTATGTATAGAA |
| | | | *Alu*-free fragment #8 | | TTCTATACATACATGTCCTAATGGGTGAAACTCTATCTCAAAAA | CTTTAATGGCAGGCTCTTTTCTTTTACCTAGTACTCTGTGTATTATGGG |
| | | | *Alu*-free fragment #9 | | CCCATAATACACAGAGTACTAGGTAAAAGAAAAGAGCCTGCCATTAAAG | CCTACTGTAAGTGTTTGTTACACCACTTTTCTGTTGTTTGCACTGA |
| | | | *Alu*-free fragment #10 | | TCAGTGCAAACAACAGAAAAGTGGTGTAACAAACACTTACAGTAGG | TCAAAGGCTTTGCCATGTTTCGCCATGAGGATGCTCTCTTTCTTTTTCT |
| | | | *Alu*-free fragment #11 | | AGAAAAAGAAAGAGAGCATCCTCATGGCGAAACATGGCAAAGCCTTTG | TTCCCAACTCCCTTCCTGTATCCCACCTCCTGTTAACAAG |
| | | | *Alu*-free fragment #12 including right HA arm | | CTTGTTAACAGGAGGTGGGATACAGGAAGGGAGTTGGGAA | CTTGCATGCAGTCGACGGGGATCCAAGTTCTGCATCCCCAGGAG |
| pTO642 | Mutating payload for creating ~2.5-kb non-*Alu* sequences deleted *TP53* 1st intron cell | HR110PA-1 | Non-*Alu* sequence deleted fragment #1 | pTO467 | GTAAAACGACGGCCAGTGAATTCTTCCCCAGAGGTATCTTCCA | TTAAAATTAAAAGCTTTTTTTTGAGACGGAGTTTCGTTCTTGT |
| | | | Non-*Alu* sequence deleted fragment #2 | pTO467 | TCTCAAAAAAAAGCTTTTAATTTTAATTATTTTACAGTTGGAG | GCTCAACAAAGGTTAGCTCTATATTGCCCTGTAACCTGCAA |
| | | | Non-*Alu* sequence deleted fragment #3 | pTO467 | AGAGCTAACCTTTGTTGAGC | TCAAAGGCTTTGCCATGTTT |
| | | | Non-*Alu* sequence deleted fragment #4 | pTO467 | AAACATGGCAAAGCCTTTGACTTGTTAACAGGAGGTGGGA | CTTGCATGCAGTCGACGGGGATCCAAGTTCTGCATCCCCAGGAG |
| pTO643 | Mutating payload for Group I *Alu* sequences deleted *TP53* 1st intron cell | HR110PA-1 | Sequence from left HA arm to downstream of group I Alu excluding | pTO504 | GTAAAACGACGGCCAGTGAATTCTTCCCCAGAGGTATCTTCCA | AGCACTGAGTTTTCTACCATTATG |
| | | | Sequence from downstream of group I *Alu* to right | pTO467 | CATAATGGTAGAAAACTCAGTGCT | CTTGCATGCAGTCGACGGGGATCCAAGTTCTGCATCCCCAGGAG |

**(Continued)**

| Plasmid | Description | Backbone | PCR target | PCR template | Forward primer | Reverse primer |
|---|---|---|---|---|---|---|
| pTO644 | Mutating payload for Group II *Alu* sequences deleted *TP53* 1st intron cell | HR110PA-1 | Sequence from left HA arm to downstream of | pTO467 | GTAAAACGACGGCCAGTGAATTCTTCCCCAGAGGTATCTTCCA | AGCACTGAGTTTTCTACCATTATG |
| | | | Sequence from downstream of group I *Alu* to downstream of group II *Alu* | pTO504 | CATAATGGTAGAAAACTCAGTGCT | CCTACAGATAGAGGGATAACT |
| | | | Sequence from downstream of group II *Alu* to right | pTO467 | AGTTATCCCTCTATCTGTAGG | CTTGCATGCAGTCGACGGGGATCCAAGTTCTGCATCCCCAGGAG |
| pTO645 | Mutating payload for Group III *Alu* sequences deleted *TP53* 1st intron cell | HR110PA-1 | Sequence from left HA arm to downstream of | pTO467 | GTAAAACGACGGCCAGTGAATTCTTCCCCAGAGGTATCTTCCA | CCTACAGATAGAGGGATAACT |
| | | | Sequence from downstream of group II *Alu* to right HA arm excluding *Alu* sequences | pTO504 | AGTTATCCCTCTATCTGTAGG | CTTGCATGCAGTCGACGGGGATCCAAGTTCTGCATCCCCAGGAG |
| pTO607 | UKiS donor (puromycin resistance gene) for *APP* locus | HR110PA-1 | *APP* left arm | HCT116 genome | GTAAAACGACGGCCAGTGAATTCAGGCACCCTTGTCAGCGCAA | GAGGTACCGAGCTCGAATTCTAAAGGACTGCTGCTAGGGT |
| | | | *APP* right arm | HCT116 genome | GCAACGCGATCGCGTAAGGGTTTATAGAATAATGTGGGAAGAAACAAAC | CTTGCATGCAGTCGACGGGGATCCCCTTCAAAGAAAAGTCTTGCC |
| pTO608 | UKiS donor (blasticidin resistance gene) for *APP* locus | HR110PA-1 | *APP* left arm | HCT116 genome | GTAAAACGACGGCCAGTGAATTCAGGCACCCTTGTCAGCGCAA | GAGGTACCGAGCTCGAATTCTAAAGGACTGCTGCTAGGGT |
| | | | *APP* right arm | HCT116 genome | GCAACGCGATCGCGTAAGGGTTTATAGAATAATGTGGGAAGAAACAAAC | CTTGCATGCAGTCGACGGGGATCCCCTTCAAAGAAAAGTCTTGCC |
| pTO619 | UKiS donor (puromycin resistance gene) for *CD44* locus | HR110PA-1 | *CD44* left arm | HCT116 genome | GTAAAACGACGGCCAGTGAATTCCCAGAAGGACATAAGGAAAG | GAGGTACCGAGCTCGAATTCACCGAACCTGGCAGAGGCTG |
| | | | *CD44* right arm | HCT116 genome | GCAACGCGATCGCGTAAGGGGAGAGGCCAGCAAGTCTCAG | CTTGCATGCAGTCGACGGGGATCCGGACCCAGACAGTGCTGGTT |
| pTO620 | UKiS donor (blasticidin resistance gene) for *CD44* locus | HR110PA-1 | *CD44* left arm | HCT116 genome | GTAAAACGACGGCCAGTGAATTCCCAGAAGGACATAAGGAAAG | GAGGTACCGAGCTCGAATTCACCGAACCTGGCAGAGGCTG |
| | | | *CD44* right arm | HCT116 genome | GCAACGCGATCGCGTAAGGGGAGAGGCCAGCAAGTCTCAG | CTTGCATGCAGTCGACGGGGATCCGGACCCAGACAGTGCTGGTT |
| pTO621 | UKiS donor (puromycin resistance gene) for *MET* locus | HR110PA-1 | *MET* left arm | HCT116 genome | GTAAAACGACGGCCAGTGAATTCGAGAGCCGGAACGAACTCAA | GAGGTACCGAGCTCGAATTCTCCGCGGGTTCCGAGGACCG |
| | | | *MET* right arm | HCT116 genome | GCAACGCGATCGCGTAAGGGTCGCAAGCAATTGGAAACAA | CTTGCATGCAGTCGACGGGGATCCCCACAATCTGTCCAGACACAT |
| pTO622 | UKiS donor (blasticidin resistance gene) for *MET* locus | HR110PA-1 | *MET* left arm | HCT116 genome | GTAAAACGACGGCCAGTGAATTCGAGAGCCGGAACGAACTCAA | GAGGTACCGAGCTCGAATTCTCCGCGGGTTCCGAGGACCG |
| | | | *MET* right arm | HCT116 genome | GCAACGCGATCGCGTAAGGGTCGCAAGCAATTGGAAACAA | CTTGCATGCAGTCGACGGGGATCCCCACAATCTGTCCAGACACAT |
| pTO625 | Mutating payload for creating *APP* all intron deleted cell | HR110PA-1 | sequence from *APP* left arm to 1st exon | HCT116 genome | GTAAAACGACGGCCAGTGAATTCAGGCACCCTTGTCAGCGCAA | TGCTGTGCGAGTGGGATC |
| | | | sequence from *APP* exon to right arm | HCT116 cDNA | GATCCCACTCGCACAGCA | CTTGCATGCAGTCGACGGGGATCCCCTTCAAAGAAAAGTCTTGCC |
| pTO631 | Mutating payload for creating *CD44* all intron deleted cell | HR110PA-1 | sequence from *CD44* left arm to 1st exon | HCT116 genome | GTAAAACGACGGCCAGTGAATTCCCAGAAGGACATAAGGAAAG | GAGGACGGGACGAGGATG |
| | | | sequence from *CD44* exon to right arm | HCT116 cDNA | CATCCTCGTCCCGTCCTC | CTTGCATGCAGTCGACGGGGATCCGGACCCAGACAGTGCTGGTT |
| pTO632 | Mutating payload for creating *MET* all intron deleted cell | HR110PA-1 | sequence from *MET* left arm to 1st exon | HCT116 genome | GTAAAACGACGGCCAGTGAATTCGAGAGCCGGAACGAACTCAA | CTCCTCTCAGCAAGTCAGCT |
| | | | sequence from *MET* exon to right arm | HCT116 cDNA | AGCTGACTTGCTGAGAGGAG | CTTGCATGCAGTCGACGGGGATCCCCACAATCTGTCCAGACACAT |
| pTO454 | *TP53*-specific gRNA/Cas9 expression plasmid for UKiS 1st step in HCT116 or iPS | pX330 | *TP53*-gRNA (R) target sequence | Oligonucleotide only | CACCGCTCAGAGGGGGCTCGACGCT | AAACAGCGTCGAGCCCCCTCTGAGC |
| pTO639 | *TP53*-specific gRNA/Cas9 expression plasmid for UKiS 1st step in iPS cells | pX330 | *TP53*-gRNA (L) target sequence | Oligonucleotide only | CACCGGTGCTTTAAGAATTACCGC | AAACGCGGTAATTCTTAAAGCACC |
| pTO546 | *APP*-specific gRNA/Cas9 expression plasmid for UKiS 1st step in HCT116 cells | pX330 | *APP*-gRNA (L) target sequence | Oligonucleotide only | CACCgCTCCCGGGGGTGTCGTATAA | AAACTTATACGACACCCCCGGGAGc |

**(Continued)**

| Plasmid | Description | Backbone | PCR target | PCR template | Forward primer | Reverse primer |
|---|---|---|---|---|---|---|
| pTO548 | *APP* -specific gRNA/Cas9 expression plasmid for UKiS 1st step in HCT116 cells | pX330 | *APP* -gRNA (R) target sequence | Oligonucleotide only | CACCgTTCTATAAATGG ACACCGAT | AAACATCGGTGTCCATT TATAGAAc |
| pTO595 | *CD44* -specific gRNA/Cas9 expression plasmid for UKiS 1st step in HCT116 cells | pX330 | *CD44* -gRNA (L) target sequence | Oligonucleotide only | CACCgCGAGGATGGCG GACCGAACC | AAACGGTTCGGTCCGC CATCCTCGc |
| pTO599 | *CD44* -specific gRNA/Cas9 expression plasmid for UKiS 1st step in HCT116 cells | pX330 | *CD44* -gRNA (R) target sequence | Oligonucleotide only | CACCGCCAAGTGGACT CAACGGAG | AAACCTCCGTTGAGTC CACTTGGC |
| pTO603 | *MET* -specific gRNA/Cas9 expression plasmid for UKiS 1st step in HCT116 cells | pX330 | *MET* -gRNA (L) target sequence | Oligonucleotide only | CACCGGGCCGCGCGC GCCGATGCC | AAACGGCATCGGCGCG CGCGGCCC |
| pTO604 | *MET* -specific gRNA/Cas9 expression plasmid for UKiS 1st step in HCT116 cells | pX330 | *MET* -gRNA (R) target sequence | Oligonucleotide only | CACCGTTCCCACCTCG CAAGCAAT | AAACATTGCTTGCGAG GTGGGAAC |
| pTS195 | Off-Target Less gRNA/Cas9 expression plasmid for UKiS 2nd step | pX330 | TL-gRNA target sequence | Oligonucleotide only | CACCGGCGCAACGCGA TCGCGTAA | AAACTTACGCGATCGC GTTGCGCC |

### Primers used for plasmid sequencing

| Target | Sequence |
|---|---|
| UKiS 1st step donor plasmids | CTGCAAGGCGATTAAGTTGG |
| | GGGCTGTCCCTCATAAAAGT |
| | ACCCGTTGCGAAAAAGAAC |
| | ACAGTCCCCGAGAAGTTGG |
| | ATGGGTACCGTGAGCAAGG |
| | AGCAGAAGAACGGCATCAAGGT |
| | GAGGGCAGAGGAAGTCTTCT |
| | CAGACATGATAAGATACATTGATG |
| | AACTTGTTTATTGCAGCTTA |
| | TCACGTAAGTAGAACATGAAATAACC |
| | CCCTTACGCGATCGCGTTGC |
| | TATGCTTCCGGCTCGTATGT |
| UKiS 2nd step donor plasmids for *TP53* 1st intron mutation | CTGCAAGGCGATTAAGTTGG |
| | CCTGGTCTGAAGGACAGCA |
| | GGTAATTCTTAAAGCACCTGCAC |
| | CTCCCCAACTCCATTTCCTT |
| | TCTCGGCTCCGTGTATTTTC |
| | CCCATGGCATCCTAGTGAAA |
| | AATCCCTCCCCTTCTTTGAA |
| | TCCTAAATCGAGGTGGCATT |
| | GGAGAACTTCATATAGAATGGAATCA |
| | GGAATTGCGAGTTTGGAAGT |
| | TTAACCCCAGGGTCATGAAG |
| | TCTTAGTGCTATTTTCTATTTATCTCC |
| | CCAATTTTAGCAAACCAAACC |
| | ACATTCTAACCAAATAATGTTTA |
| | CATAATGGTAGAAAACTCAGTGCT |
| | CGCTTTTTCCTCTAAGATCAGG |
| | CCCATTAGGACATGTATGTATAGAA |
| | AATAACGATTAGAAAGGAAGAAATA |
| | AATAACGATTAGAAAGGAAGAAATA |
| | CCCATTAGGACATGTATGTATAGAA |
| | AGTGAAACTCTATCTCAAAAA |
| | AGGGCACCTGCATTTCTCTT |
| | TACTGTGTGCCCAGCACTTT |
| | GCTGTTTGGTAGGGGATGTT |
| | GTGTAACAAACACTTACAGTAG |
| | AAACATGGCAAAGCCTTTGA |
| | GAATAACACACAAGCCTGTTATA |
| | AGGTGAGCGGTGAGACAGTT |
| | ACAGGAAGGGAGTTGGGAATA |
| | CTTCCTGAAAACAACGTTCTG |
| | TATGCTTCCGGCTCGTATGT |
| UKiS 2nd step donor plasmid for *APP* all introns deletion | CTGCAAGGCGATTAAGTTGG |
| | AGCGGTAGGCGAGAGCAC |
| | AACCCCAGATTGCCATGTT |
| | CCACTGGCTGAAGAAAGTGAC |
| | TTGCCCGAGATCCTGTTAAA |
| | TGACACACCTCCGTGTGATT |
| | TGCAGAAGATGTGGGTTCAA |
| | TGCATGAATAGATTCTCTCCTGA |
| | TGCTGGTCTTCAATTACCAAGA |
| | TATGCTTCCGGCTCGTATGT |

**(Continued)**

| Target | Sequence |
|---|---|
| UKiS 2nd step donor plasmid for *CD44* all introns deletion | CTGCAAGGCGATTAAGTTGG |
| | CCAGCCTCTGCCAGGTTC |
| | CCACGTGGAGAAAAATGGTC |
| | CCATCCCAGACGAAGACAGT |
| | CCATGAGCATCATGAGGAAG |
| | GGAATGATGTCACAGGTGGA |
| | GAGAGGCCAGCAAGTCTCAG |
| | CAGGGTTAATAGGGCCTGGT |
| | TATGCTTCCGGCTCGTATGT |
| UKiS 2nd step donor plasmid for *MET* all introns deletion | CTGCAAGGCGATTAAGTTGG |
| | GAGCGCCTCAGTCTGGTC |
| | CGCCGTGATGAATATCGAAC |
| | GAGGGACAAGGCTGACCATA |
| | TTTATTAGTGGTGGGAGCACAA |
| | TTCAACCGTCCTTGGAAAAG |
| | AAGAGGGCATTTTGGTTGTG |
| | AGTGAAGTGGATGGCTTTGG |
| | GGCCATCGATATTCTTTGCT |
| | GGCTGTAGTGCAGTGGTGTG |
| | CAGCATGTTTGTAAAGCAGGA |
| | GAAAAGGTATGTCAGACTGGGATT |
| | TATGCTTCCGGCTCGTATGT |
| gRNA/Cas9 expression plasmids | GGAAAGTCCCTATTGGCGTTA |

### Supplementary Table 5. Primers used for off-target analysis

#### Primers used for PCR

| Fig. # | Target | Forward primer | Reverse primer |
|---|---|---|---|
| Supplementary Fig. 2 | *TP53*- specific gRNA off-target candidate #1 | ACAGAGGCCCCCAAATAAG | TGACTGTGTCTGGAGGATGG |
| | *TP53*- specific gRNA off-target candidate #2 | GAACCTCCAGTGTGTGAAAGG | ATACAAGAAATGCCCCCAAA |
| Supplementary Fig. 3 | TL-gRNA off-target candidate | CGACCTTCTGGGAGCAGT | TCCACAGGTACGACAACGAA |

#### Primers used for sequencing

| Fig. # | Target | primer |
|---|---|---|
| Supplementary Fig. 2 | *TP53*- specific gRNA off-target candidate #1 | ACAGAGGCCCCCAAATAAG |
| | *TP53*- specific gRNA off-target candidate #2 | GAACCTCCAGTGTGTGAAAGG |
| Supplementary Fig. 3 | TL-gRNA off-target candidate | CGACCTTCTGGGAGCAGT |

### Supplementary Table 6. Primers used for real-time RT-PCR

| Fig. # | Target | Forward primer | Reverse primer |
|---|---|---|---|
| Fig. 5c and Fig. 6b-c | *TP53* mRNA for qRT-PCR | GACTGCCTTCCGGGTCACT | GGACAGCATCAAATCATCCA |
| | 18S rRNA for qRT-PCR | ATTAATCAAGAACGAAAGTCGGAG | TTTAAGTTTCAGCTTTGCAACCAT |
| Supplementary Fig. 8 | *CDKN1A* mRNA for qRT-PCR | AGGGGACAGCAGAGGAAGAC | GGCGTTTGGAGTGGTAGAAA |
| | *GADD45A* mRNA for qRT-PCR | GGCCCGGAGATAGATGACTT | GTTTTCCTTCCTGCATGGTT |
| | *BAX* mRNA for qRT-PCR | CATGTTTTCTGACGGCAACT | CCGGAGGAAGTCCAATGTC |
| | *PUMA* mRNA for qRT-PCR | TCAGGAAAGGCTGTTGTGCT | ACGTTTGGCTCATTTGCTCT |
| | *RRM2B* mRNA for qRT-PCR | GCCAGGACTCACTTTTTCCA | CAATGATCTCCCTGACCCTTT |
| | *14-3-3σ* mRNA for qRT-PCR | GTTGAGCGCACCTAACCACT | CTGTCCAGTTCTCAGCCACA |
